# Lipidic and senescent macrophages predict progression and response to combinatorial immunotherapy in triple-negative breast cancer

**DOI:** 10.1101/2024.06.24.600550

**Authors:** Chun Lai Chan, Alex To, Shihui Zhang, Jason Wing Hon Wong, Yuanhua Huang, Yiming Chao, Ryohichi Sugimura

## Abstract

Immune cell subsets in the tumor predict prognosis. Identifying reliable subsets consistently in multiple patients is clinically important. What’s more advantageous to the field is if such subsets can be a target of immunotherapy. Here we analyzed single-cell RNA-sequencing datasets of patients with triple-negative breast cancer and identified APOE or FABP5-expressing macrophages correlated with poor prognosis. Further validation with TCGA-BRCA cohort detailed molecular signatures of these macrophages as lipidic and senescent. Our receptor-ligand mapping identified lipidic and senescent macrophages both suppress T-lymphoid and myeloid immunogenicity. Finally, we discovered that immune checkpoint therapy combined with chemotherapy reprogrammed anti-tumor microenvironment enriched with FOLR2^+^ macrophages facilitated T-cell activations. This suggests that lipidic and senescent macrophages could be a therapeutic target of immune checkpoint therapy.

## Introduction

Breast cancer is clinically categorized into various subtypes based on the expression of hormone receptors: estrogen receptor positive (ER^+^), progesterone receptor positive (PR^+^), and human epidermal growth factor receptor 2 positive (HER2^+^) breast cancer (Pourzand et al., 2011; Akshata Desai, 2012; Supplementary Figure S1a). Breast cancer that lacks the expression of these three hormone receptors is classified as triple-negative breast cancer (TNBC) (Akshata Desai, 2012; Supplementary Figure S1a).TNBC is the most immunosuppressive and aggressive clinical subtype of breast cancer, characterized by a high chance of metastasis and a fast growth rate (Sun, 2021; Chen et al., 2022). Around half of the individuals diagnosed with early-stage TNBC (Stage I-III) experience disease recurrence (Costa & Gradishar, 2017). Thus, it is of great clinical significance to understand the potential molecular mechanisms in aggressiveness of TNBC, which may eventually pave the way for identifying therapeutic targets.

Tumor-immune microenvironment (TIME) represents the complex ecosystem of immune and stromal cells associated with the tumor (Chew & Abastado, 2012). Vascular systems, cancer-associated fibroblasts (CAFs), myofibroblasts, cytokines, and the associated extracellular matrix (ECM) network also play an important role in regulating immune systems (Weber & Kuo, 2012). Analysis of TIME offers invaluable clinical information about the anti-tumor immune response, mechanisms of immune evasion, prevalence of pro-tumor factors, and response to anti-cancer treatments like immune-checkpoint blockade (ICB) (Binnewies et al., 2018; Kim & Cho, 2022). Due to advancements in single-cell sequencing technologies, the characterization of TIME is no longer limited to bulk resolution as it was in the past. Significant efforts have been made in recent years to unveil the single-cell and spatial transcriptome of TNBC (Wu et al., 2021; Pal et al., 2021; Zhang et al., 2021; Bassiouni et al., 2023). These efforts have enabled the identification of subtypes of immune and stromal subsets that correspond to poor survival outcomes or even poor response to immunotherapy in TNBC patients (Zhang et al., 2021).

Despite the earlier effort spent on TIME, the molecular mechanisms explaining the relationship between the immuno-suppressive TIME, tumor progression, and therapeutic response in TNBC patients have not been completely deciphered. Hence, in this study, we re-analyzed sc-RNA seq data of invasive ductal carcinoma from 19 treatment-naïve patients of TNBC (1). By integrating bulk RNA-seq data from the BRCA-TCGA cohort (ref), we identified lipidic and senescent macrophages in TNBC patients. These macrophages, featured with either FABP5 or APOE expression, predict poor patient outcomes. In line with observations in other solid tumors, collagen-rich CAF induces senescence-associated secretory phenotype (SASP) in macrophages, altogether participating in T-lymphoid and myeloid immune suppression. Intriguingly, immune checkpoint therapy in combination with chemotherapy reprogrammed TIME enriched with anti-tumor FOLR2^+^ macrophages. These indicate that lipidic and senescent macrophages could be a therapeutic target of immunotherapy in TNBC.

## Results

### FABP5+ and APOE+ macrophages predict poor prognosis in TNBC

We defined immune cell subsets that predict the prognosis of TNBC (Figure 1a-1d). We retained 66,834 cells from 19 treatment-naïve patients quality control steps. We categorized clusters of cells on Uniform Manifold Approximation and Projection (UMAP based on i. patient identity (Supplementary Figure 1f) and ii. clinical (immunohistochemical) subtypes of breast cancer (Figure 1c). We clustered into immune cells, stromal cells, and epithelial cells using canonical markers according to Wu et al. (2021) (Figure 1b and 1c). As for the immune cell clusters, we identified T/ NKT / NK cells, myeloid cells, B-cells (Supplementary Figure S3d-S3f) and the antibody-producing plasmablasts by the upregulated expression of pan T-cell marker *CD3E*, myeloid lineage marker *CD68*, *MS4A1* and *JCHAIN* respectively (Figure 1b). We then identified several types of stromal cells, including cancer-associated fibroblast (CAFs), myofibroblast, and endothelial cells. CAF was detected by *COL1A1* while endothelial cell was detected by *PECAM1* (Figure 1b). We identified myofibroblasts based on the upregulation of *ACTA2* (Figure 1b). Lastly, we annotated epithelial cells by *EPCAM* (Figure 1b). The hallmarks of breast cancer is the presence of aneuploid genome (Elenbaas et al., 2001; Pfister et al., 2018). Our copy number variations (CNVs) analysis of *EPCAM*^+^ cells distinguished the aneuploid tumorous epithelial cells from the diploid epithelial cells (Figure 4d, 4e and Supplementary Figure S4c).

**Figure 1.**
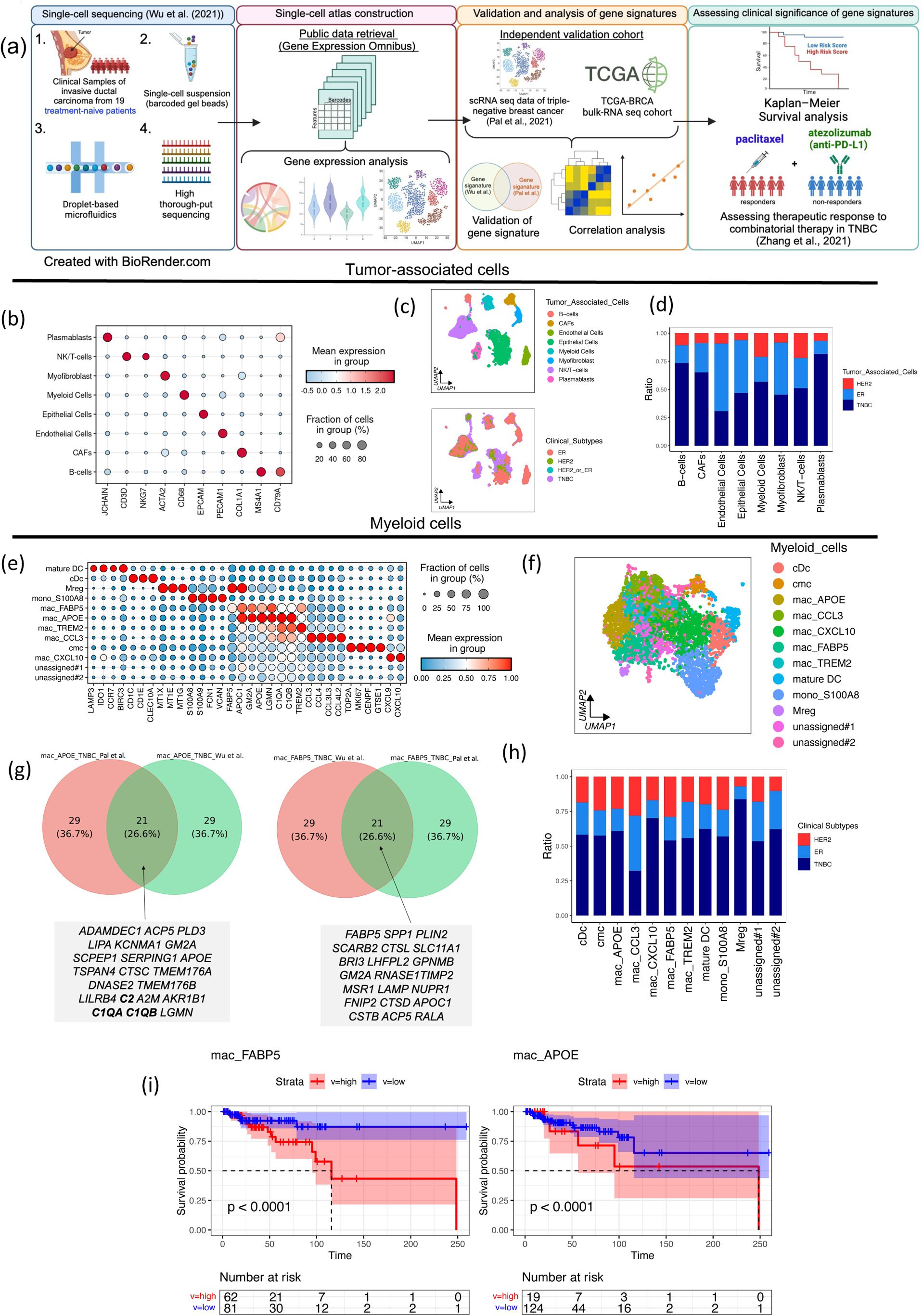
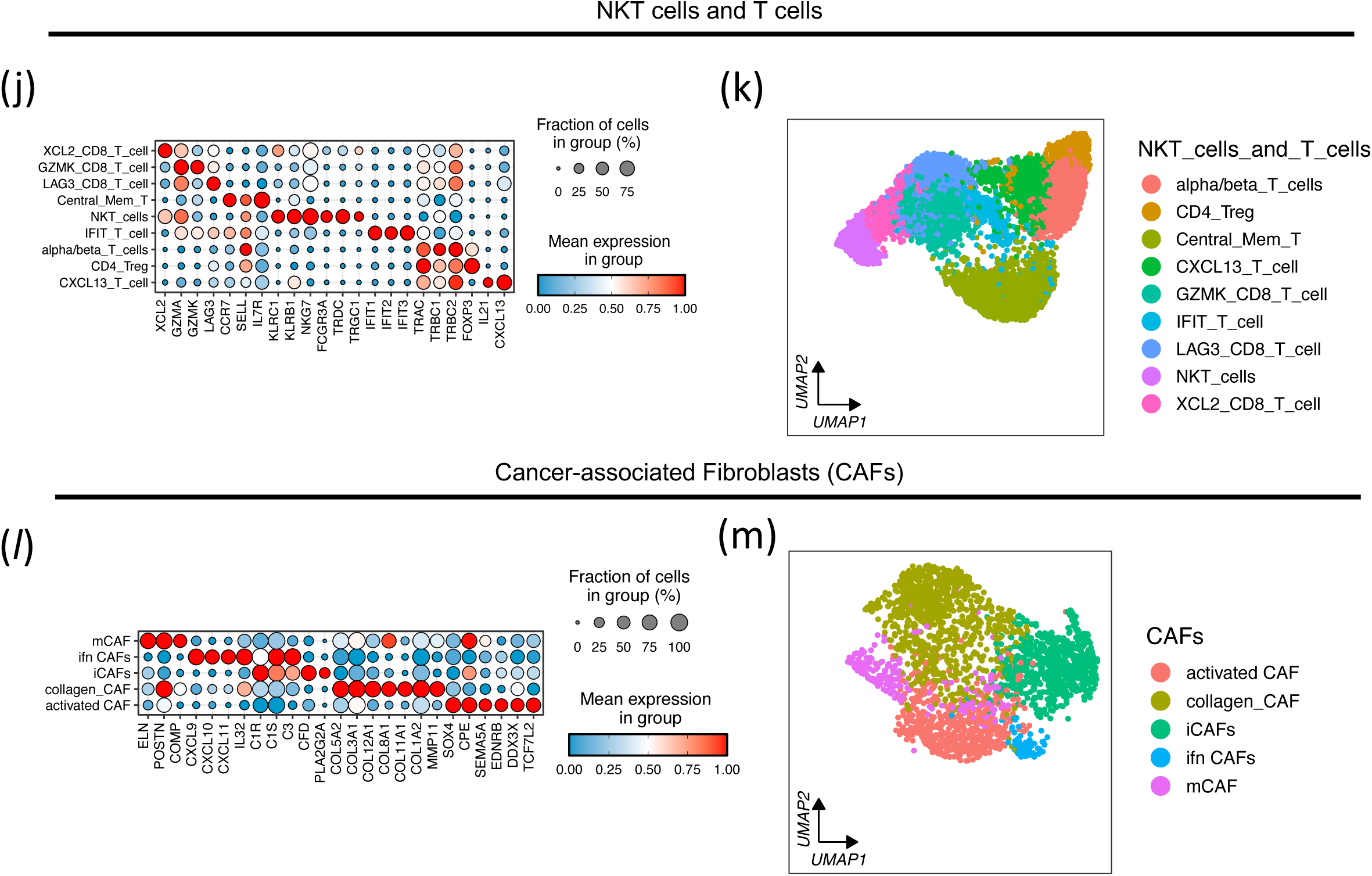
Overall Transcriptomic landscape of invasive ductal carcinoma (IDC) at single-cell resolution: (a) Schematic diagram illustrating the overall study design of this study. Created with BioRender.com (b) Bubble plot illustrating the expression of canonical markers used to categorize the tumor-associated cells in IDC. Markers were obtained from Wu et al. (2021) (c) Uniform Manifold Approximation and Projection (UMAP) plots showing clusters of cells derived from 19 patients. Cells are catergorised based on i. identity of tumor-associated cells and ii. clinical subtypes of breast cancer (d) Stacked bar chart showing the relative proportion of different types tumour-associated cells in breast cancer. (e) Bubble plot illustrating the marker genes used for identifying subtypes of tumor-associated myeloid cells. The size of each bubble is proportional to the percentages of cells expressing selected marker genes. (f) UMAP showing the clusters of tumor-associated myeloid cells (g) Venn diagrams illustrating the lipid macrophage signatures co-expressed in both sc-RNA seq datasets from Wu et al. (2021) and Pal et al. (2021). The numbers and percentage of overlapping / non-overlapping genes between the lipid-associated macrophages in Wu et al. (2021) and Pal et al. (2021) are shown on the Venn diagram. (h) Stacked bar chart showing the relative proportion of different types tumour-associated myeloid cells in breast cancer. (i) Kaplan-Meier survival analysis of the marker genes of TNBC-derived lipid-associated macrophages (mac_APOE and mac_FABP5) identified by *COSG*. The dashed lines indicate the median survival time for the high-risk and low-risk cohorts. The 95% confidence interval bands for the high-risk and low-risk cohorts are shaded in low-opacity blue and low-opacity yellow, respectively. The rectangle box below the curves contains count of individuals who are at risk over the given period. (j,l) Bubble plots illustrating the marker genes used for identifying subtypes of T-cells/ NKT cells in IDC. The size of each bubble is proportional to the percentages of cells expressing selected marker genes. (k,m) UMAP plots showing the clusters of tumor-associated T-cells/ NKT cells and CAFs in breast cancer

We identified ten subclusters of tumor-associated myeloid cells after further sub-clustering of CD68^+^ cells (Figure 1e and 1f). We first annotated mac_TREM2 and mac_CCL3 (Figure 1f), based on the expression of *TREM2*, and *CCL3* (Figure 1e), respectively. This is consistent with TREM2+ macrophages in colon cancer by Zhang et al. (2020). Then, we identified a cluster of regulatory macrophages (Mreg) (Figure 1f), as evidenced by its upregulation of metallothionein genes (i.e. *MT1M, MT1H, MT1G, MT1E, MT1F, MT1X)* (Figure 1e) reported by Gurvich et al. (2020). We identified a cluster of tumor-associated conventional dendritic cells (cDc) (Figure 1f), based on the elevated expression of dendritic cell markers *CLEC10A* and *CD1C* (Figure 1e) reported by Heger et al. (2018). We detected the monocyte S100A8_mono based on the expression of monocyte markers *FCN1*, *S100A8*, and *S100A9*. Both *VCAN* and *THBS1* were upregulated in S100A8_mono, manifesting the immunosuppressive traits of tumor-associated macrophages (TAMs) (Zhang et al., 2021) (Figure 1f). In addition, we identified cycling myeloid cells (cmc) by the expression of *MKl67* (Figure 1e) indicating cellular proliferation (Bullwinkel et al., 2005). Gene Ontology (GO) analysis revealed that cmc is enriched with biological functions related to cell cycle and cell division, such as "chromosome segregation", "mitotic sister chromatid segregation" and "DNA replication" (Supplementary Figure S1*l*).

In order to define macrophage subsets predicting patient outcomes, we identified two clusters of lipid-associated macrophages, namely mac_FABP5 and mac_APOE, based on the expression of genes essential for lipid metabolism (e.g. FABP5 and APOE) (Figure 1g). This is consistent with the observations by Wu et al. (2021). mac_FABP5 and mac_APOE constitute a greater proportion in TNBC tumours (Figure 1h) and are associated with poor survival (Figure 1i).

We then identified subtypes of tumor-associated NK/ NKT and T-cells by sub-clustering the CD3E^+^ and NKG7^+^ cells (Figure 1j, k, Supplementary Figure S3a). First, we annotated subtypes of tumor-associated CD8^+^ T-cells, namely XCL2_CD8_T_cell, GZMK_CD8_T_cell and LAG3_CD8_T_cell (Figure 1k), characterized by high expression of *XCL2, GZMK* and *LAG3*, respectively (Figure 1j). Furthermore, we identified clusters of T helper (T_H_) cells, including regulatory T-cells (T-reg) and follicular TH (T_FH_) cells (CXCL13_T_cells). While we characterized T-reg by the expression of *FOXP3* transcription factor, the CXCL13_T_cells elevated expression of *PDCD1*, *IL21* and *CXCL13* (Figure 1j), which were T_FH_ markers. We also identified a cluster of T-cells having high expression of genes related to responses to interferon, including *IFIT1*, *IFIT2* and *IFIT3* (Figure 1j), as IFIT_T_cell, and another cluster of T-cells having high expression of *CCR7* and *IL7R* as central memory T-cells (Central_Mem_T_cells) (Figure 1j). Both *CCR7* and *IL7R* were reported to be the markers for Central_Mem_T_cell (Raphael et al., 2020). Lastly, we identified a cluster of natural killer T (NKT) cells (Figure 1k), and its presence is evidenced by the elevated expression of natural killer (NK) cell markers such as *KLRC1*, *KLRB1*, *NKG7* and *FCGR3A* (encoding FC receptor *CD16*), as well as genes encoding γδ subunits of T-cell receptors (*TRDG* and *TRGC1*) (Figure 1j).

Lastly, we detected CAFs based on the single-cell transcriptomic data of the 19 patients (Supplementary Figure S3b-S3c). We adopted markers from Cords et al. (2023) to identify CAF subtypes. We identified collagen CAFs and matrix CAFs (mCAFs) (Figure 1m) based on their respective expressions of extracellular matrix (ECM) proteins (Figure 1l). While both mCAFs and collagen CAFs highly express collagen genes and are enriched in biological processes pertinent to collagen production (Figure 1l, Supplementary Figure S1j), we found that mCAFs differentially express elastin (Figure 1l), another ECM protein encoded by *ELN* (Figure 1l). Additionally, mCAFs are enriched in GO terms like "extracellular matrix organization", "extracellular structure organization" and "heparin-binding" (Supplementary Figure S1k), suggesting mCAFs in remodeling the ECM of the tumor immune microenvironment (TIME) of breast cancer. Moreover, mCAFs interact with tumor-associated T-cells through TGF-β signaling, a known immune-suppressor (Supplementary Figure S2h). We then identified another CAF subtype, namely inflammatory CAFs (iCAFs) (Figure 1m), based on their upregulated expression of complement proteins (*C3*, *CFD*) (Figure 1l). Lastly, we annotated the interferon-response CAFs (Figure 1m) based on their expression of chemokines *CXCL9*, *CXCL10* and *CXCL11* (Figure 1l), which are the three chemokines involved in the interferon response as well as *IL32* (Figure 1l), a marker for chronic inflammatory response.

### Lipid TAMs are enriched in senescence-associated secretory phenotype (SASP) signature

We focused mac_FABP5 and mac_APOE as they predict patient outcomes. We linked their molecular signatures with the inflammatory status (Figures 2a-f). The Spearman correlation analysis using the TCGA-BRCA cohort has revealed that both mac_APOE and mac_FABP5 exhibited a strong correlation with the inflammatory signature reported adapted from Zhang et al. (2021) (Figure 2a and 2d). Notably, the co-expression of lipid-associated macrophages and inflammatory signatures correspond to the poor survival probability (Figure 2b and 2e). The GO analysis also revealed that both mac_APOE and mac_FABP5 enriched biological processes related to lipid metabolism, such as ‘lipid transport’, ‘regulation of lipid localization’, ‘lipoprotein catabolic process (Figure 2c and 2f). mac_APOE in TNBC (mac_APOE_TNBC) enriched genes related to the activation of innate-immunity-related pathways such as compliment pathway (Figure 2d and 2f), suggesting lipidic macrophages in inducing inflammation in TNBC. To solve why poor prognosis correlated with inflammatory signature, we dissected their composition. Both mac_APOE and mac_FABP5 express high levels of senescent-associated secretory phenotype (SASP) proteins (Figure 2g). SASP is a secretome generated by senescent cells and counteracts cytotoxic T-cells. (Prieto et al., 2023). Altogether, the lipid-associated macrophages manifest the SASP signature suppressive to immunogenicity.

**Figure 2.**
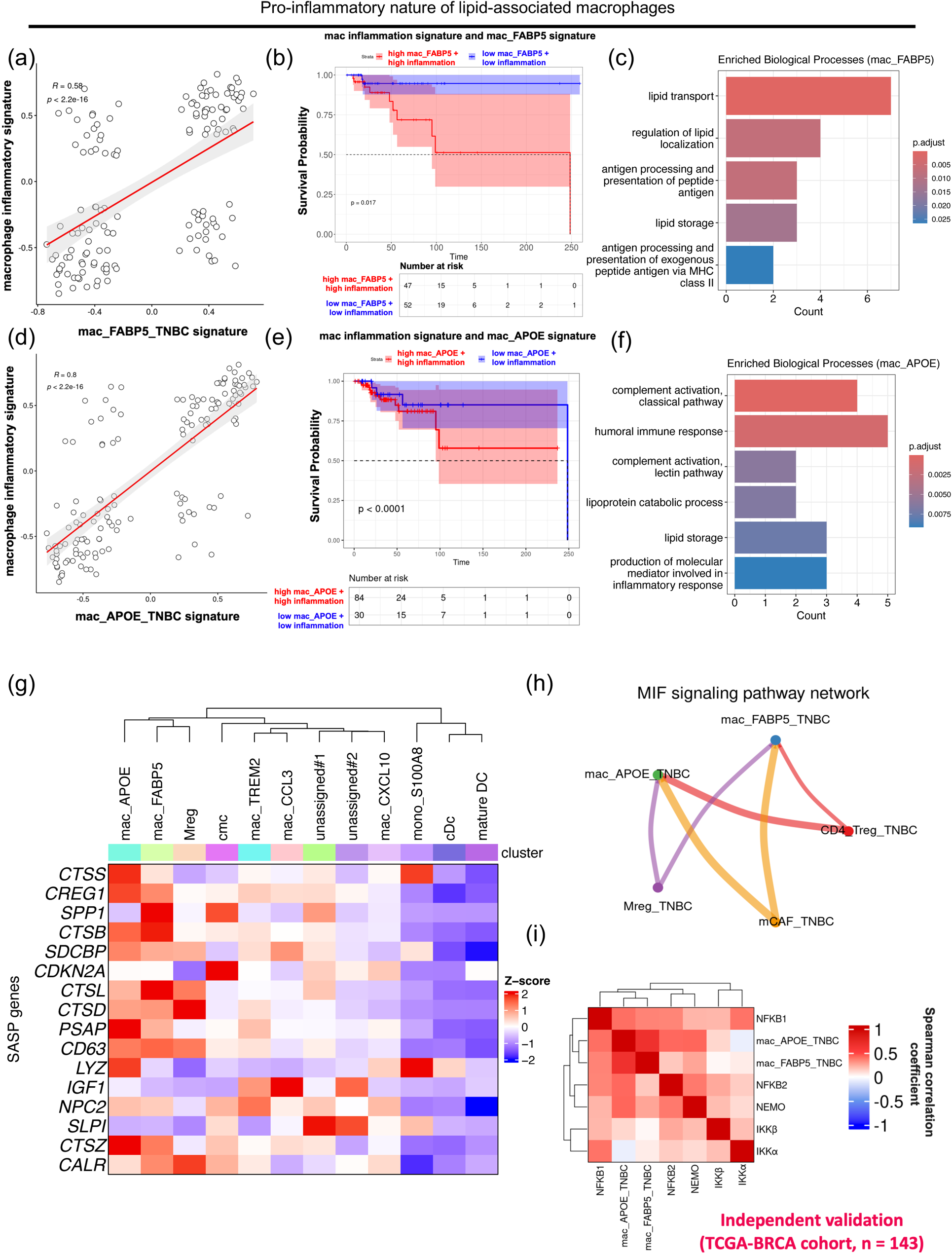
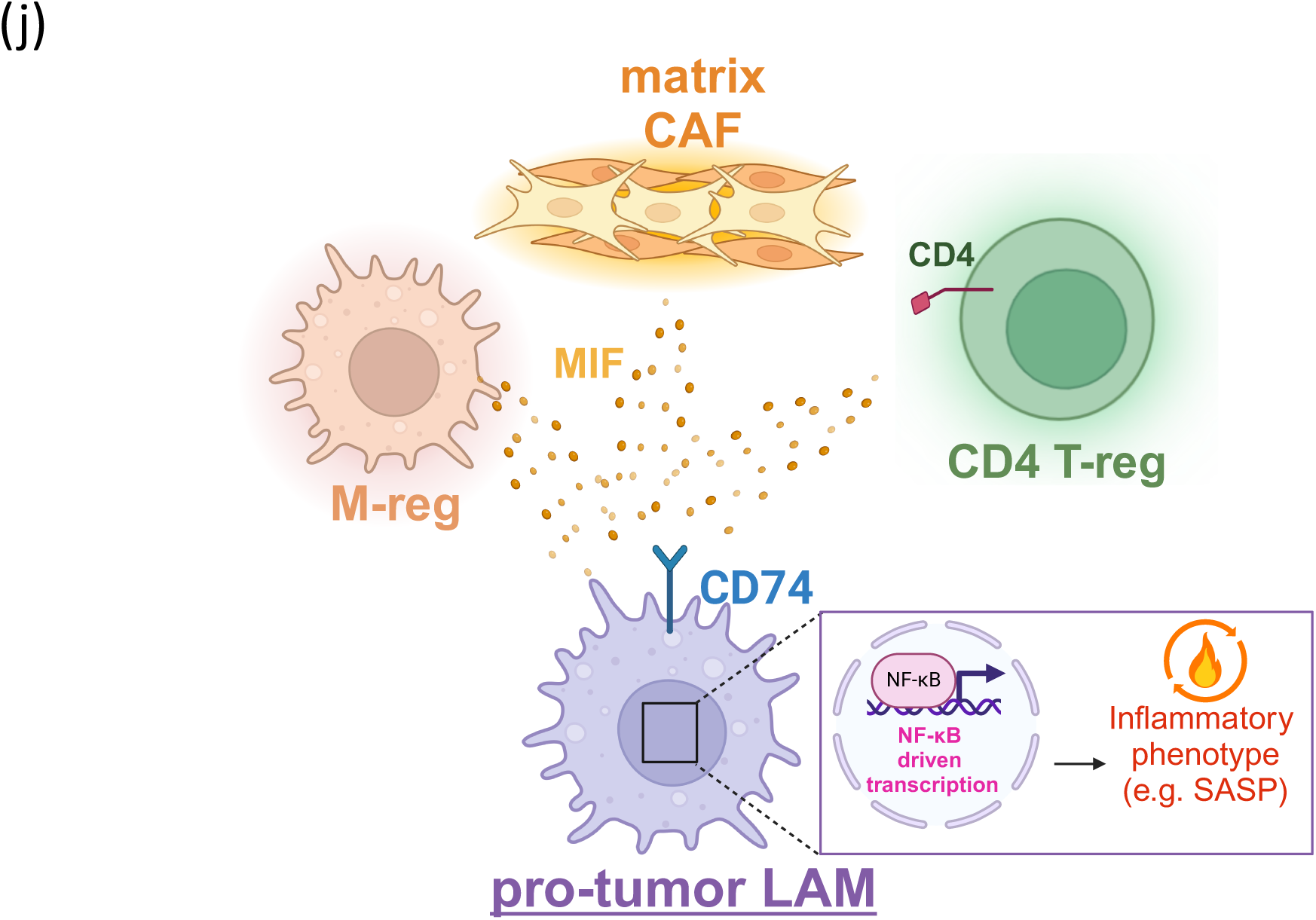
Lipid-associated macrophages exhibit a pro-inflammatory phenotype in TNBC: (a,d) Spearman correlation analysis between the myeloid inflammatory signature and the gene signatures of mac_FABP5/mac_APOE in TNBC. The correlation analysis was conducted using the bulk RNA seq data from the basal-like breast cancer patients from the TCGA-BRCA cohort. (b,e) Kaplan-Meier survival analysis of basal-like breast cancer patients from the TCGA-BRCA cohort expressing high or low levels of macrophage inflammatory signatures. The analysis compared patients who co-expressed high levels of the macrophage inflammatory signature and the mac_FABP5/mac_APOE signature (red line) to those who expressed low levels of both signatures (blue line). (f,c) Gene ontology analysis of the gene signatures expressed by mac_APOE and those of mac_FABP5 in TNBC. (g) Heatmap illustrating the expression of genes encoding senescence-associated secretory phenotype (SASP) proteins in the myeloid subtypes. Genes with positive Z-scores are assigned with red color while those with negative scores are assigned with purple-blue color. (h) Cell-cell communication network between lipid-associated macrophages and other subtypes of tumor-associated cells that provide MIF signaling. The Communication network is identified using *CellChat* R package. (i) Heatmap showing the Spearman correlation between the enrichment score of genes associated with NF-κB pathway and the lipid macrophage signature. The Spearman correlation coefficient (R) ranges from 0 to 1 in this heatmap. Colors of the heatmap correspond to the magnitude of R (red: 0 < R < 1; white: R = 0; blue:-1 < R < 0) (j) Schematic diagram illustrating the regulation of inflammatory program of LAM by CD4 T-reg, M-reg and matrix CAF. Created with BioRender.com

We next sought to determine what signals induce SASP expression in lipid tumor-associated macrophages (TAMs). Our cell-cell communication pattern using *CellChat* (Jin et al., 2021; Supplementary Figure S2a-S2g) demonstrated that Macrophage migration inhibitory factor (MIF) signaling as the top contributor to the secreted signaling in lipid macrophages (Supplementary Figure S1f). The lipid macrophages were regulated by MIF signaling, specifically the MIF-(CD74-CD44) axis, from mCAF_TNBC, Treg_TNBC and Mreg_TNBC (Figure 2h and 2j). In concordance with Calandra and Roger (2003), MIF signaling suppresses the glucocorticoid-dependent expression of the inhibitor of nuclear factor-κB (IκB), thereby facilitating the expression of inflammatory genes through NF-κB. MIF signaling drives the inflammatory lipidic TAM. Our independent validation using bulk RNA-seq data of the TCGA-BRCA cohort revealed that both mac_FABP5_TNBC and mac_APOE_TNBC exhibit modest positive correlations with the expression of *NFKB1* and *NFKB2* (Figure 2i), which encode the p45 and p100 subunits of NF-κB, respectively. Moreover, we found that both mac_FABP5_TNBC and mac_APOE_TNBC were positively associated with the expression of IKK-β, a subunit of the IKK complex, and NEMO, the regulatory subunit of the IKK complex (Figure 2i), and importantly the IKK complex is involved in the activation of the NF-κB pathway by promoting the degradation of IκB. In line with Faget et al. (2019), inflammatory SASP of lipid macrophages is induced by MIF signaling.

### Lipid TAMs are lymphoid-suppressive via HLA mechanisms

Given the lipid-associated TAMs secreting SASP indicate T-cell suppression, we sought its molecular mechanisms. Lipid-associated TAMs could exhaust T-cells through persistent antigen presentation (Figure 3l). The effector CD8^+^ T-cells associated with the TNBC, namely the LAG3_CD8_T_cell, GZMB_CD8_T_cell and XCL2_CD8_Tcell, express higher levels of genes encoding T-cell immune checkpoints, including *PDCD1* (encoding PD-1), *TIGIT*, *HAVCR2* (encoding TIM-3), *LAYN* and *LAG3*, comparing with those in ER^+^ and HER2^+^ breast cancer (Figure 3a and 3c). Furthermore, these effector T-cell subsets have lower expression of genes encoding T-cell effector molecules, including *IFNG*, *IL2* and *TNF* (Figure 3b and 3d). Notably, these subsets of effector T-cells in TNBC downregulate the expression of *CD44* (Figure 3b), which is the activation marker during the transition from naïve T-cells to effector T-cells. These immunosuppressive T-cells cluster with each other (Figure 3a and 3b), suggesting a tendency for CD8^+^ T-cells to manifest immunosuppressive phenotype in TNBC.

**Figure 3.**
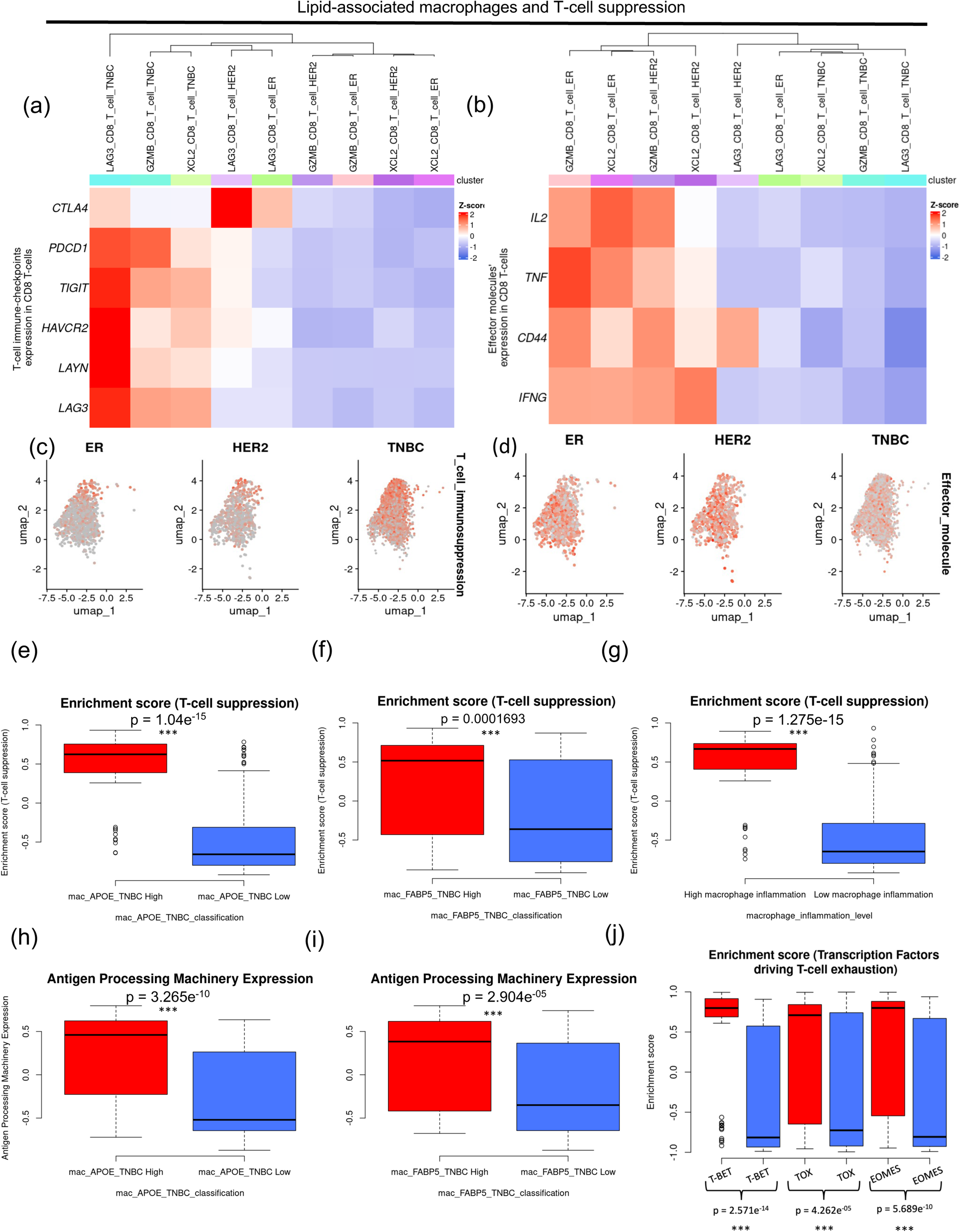
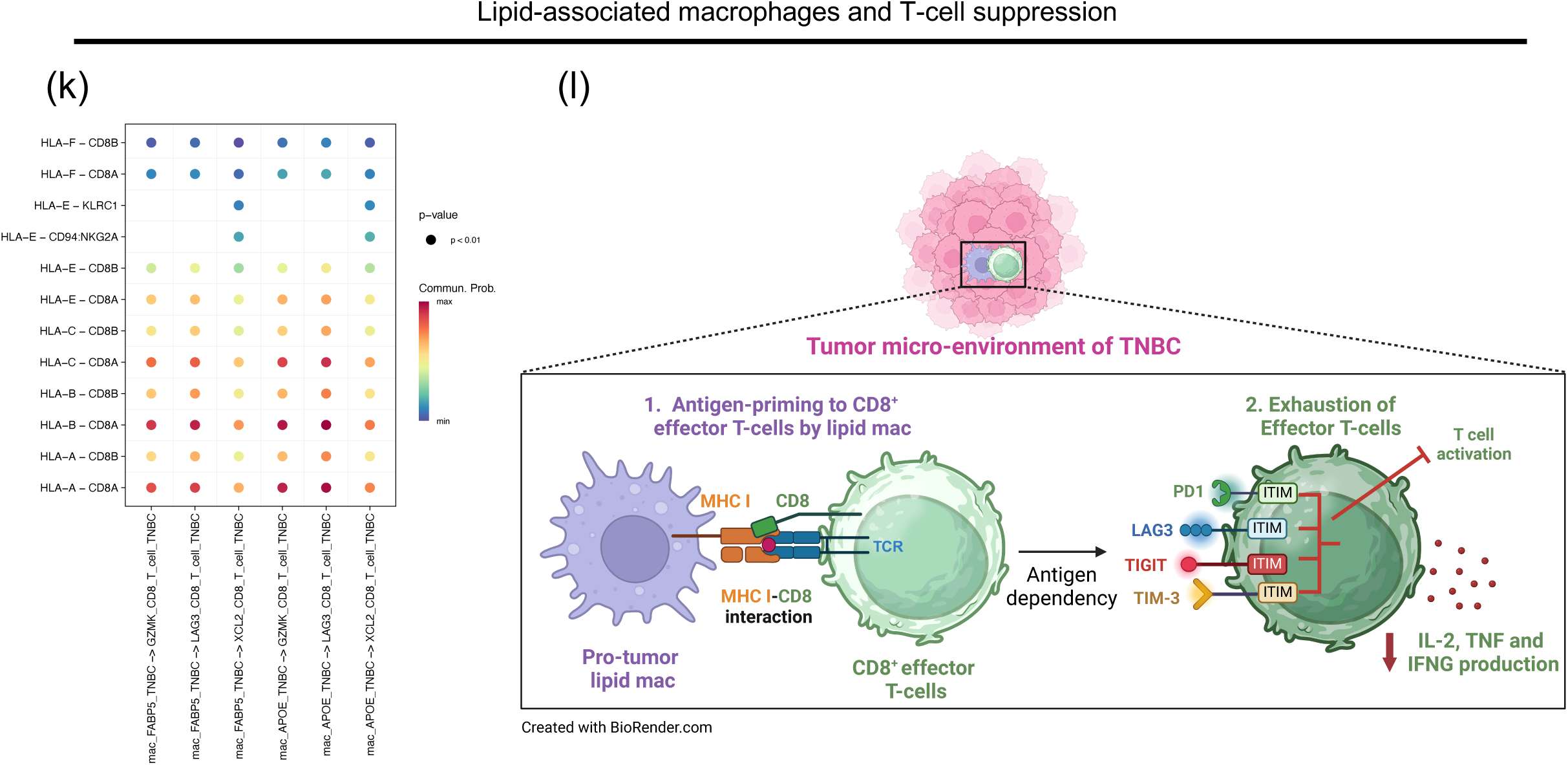
Lipid TAMs are associated with lymphoid-suppressive features in the TIME of TNBC. (a-d) Transcriptional dynamics of subtypes of effector T-cells: The marker genes used to decipher the transcriptional dynamics of subtypes of effector T-cells were adopted from Baaten et al. (2015) (54). The genes used for the analysis of transcriptional dynamics included those (a) encoding immune checkpoints and (b) effector molecules of CD8^+^ T-cells. The associated feature plots depicted (c) higher expression of immune checkpoint molecules and (d) lower expression of effector molecules in the effector CD8^+^ T-cells of TNBC. (e-g) Boxplots illustrating the distribution of T-cell suppression scores across basal-like breast cancer patients from the TCGA-BRCA cohort, showing differences based on expression of the mac_APOE_TNBC and macrophage inflammation signatures. (g) The boxplot for antigen processing machinery expression between basal-like breast cancer patients with high or low mac_APOE_TNBC signature is also shown. The median is represented by the midline, and the whiskers denote 1.5 x interquartile range. (h) Bubble plot illustrating the cell-cell communication pattern between mac_APOE_TNBC and the CD8^+^ T-cells via MHC I-CD8 axis. The associated boxplot illustrates the enrichment score of antigen processing machinery across basal-like breast cancer patients having higher or lower expression of mac_APOE_TNBC signature. (i) Schematic diagram illustrating the potential relationship between the pro-tumor lipid-associated macrophage and the cytotoxic CD8^+^ T-cells in TNBC. Created with BioRender.com.

Lipid-associated macrophages express immunosuppressive checkpoints in TNBC. We explored the relationship between lipid macrophages and the exhaustion phenotype of CD8^+^ T-cells in the bulk RNA sequencing data from the TCGA-BRCA cohort. Basal-like breast cancer patients with higher expression of the mac_APOE_TNBC/ mac_FABP5_TNBC signature exhibit a higher T-cell suppression score, compared to those with lower expression of the mac_APOE_TNBC/ mac_FABP5_TNBC signature (Figure 3e and 3f). High expression of the macrophage inflammatory signature adapted from Zhang et al. (2021) corresponds to higher expression of T-cell suppression score (Figure 3g). Additionally, higher expression of the mac_APOE_TNBC/ mac_FABP5_TNBC signature was accompanied by a significantly higher expression level of transcription factors driving T-cell exhaustion, including T-BET, TOX, and EOMES (Figure 3j). It is worth noting that T-BET and EOMES are transcription factors highly upregulated in transitory exhausted T-cells and terminal exhausted T-cells, respectively (Franco et al., 2020; Belk et al., 2022). Our cell-cell communication analysis using *CellChat* revealed strong enrichment in contact-dependent signaling between CD8^+^ T-cell subsets and mac_APOE_TNBC/ mac_FABP5_TNBC via HLA-CD8 interactions (Figure 3k). The mac_APOE_TNBC/ mac_FABP5_TNBC expression level positively correlates with the expression of antigen-processing machinery reported by Thompson et al. (2020) (i.e. *RFX5, B2M, PSME1, NLRC5, and PSMB9*) (Figure 3h and 3i). These findings suggest that these CD8^+^ T-cell subsets are subjected to antigen presentation by lipid macrophages. Importantly, persistent stimulation of T-cells by antigens is one of the hallmarks of T-cell exhaustion (Wherry & Kurachi, 2015).

### Lipid TAM and aneuploid cancer cells form a myeloid suppressive environment

We next sought to determine whether myeloid immunosuppression is linked to the lipidic and senescent macrophages (Wang & Dubio, 2015). Spearman correlation showed a strong correlation between mac_APOE_TNBC signature and the expression of certain myeloid immune checkpoints, including *LILRB2*, PD1 (*PDCD1*), and *SIGLEC10* (Figure 4a). These myeloid immune checkpoints, which contain immunoreceptor tyrosine-based inhibitory motif (ITIM) in cytosol, can relay "don’t eat me" signal that dampens the anti-tumor phagocytic activity of TAMs (Bradley, 2019; Li et al., 2023; Liu et al., 2022). We found a significant association between low survival probability in patients with basal-like breast cancer and high expression of *SIGLEC10* (Figure 4b) and *LILRB2* (Supplementary Figure S4a).

**Figure 4.**
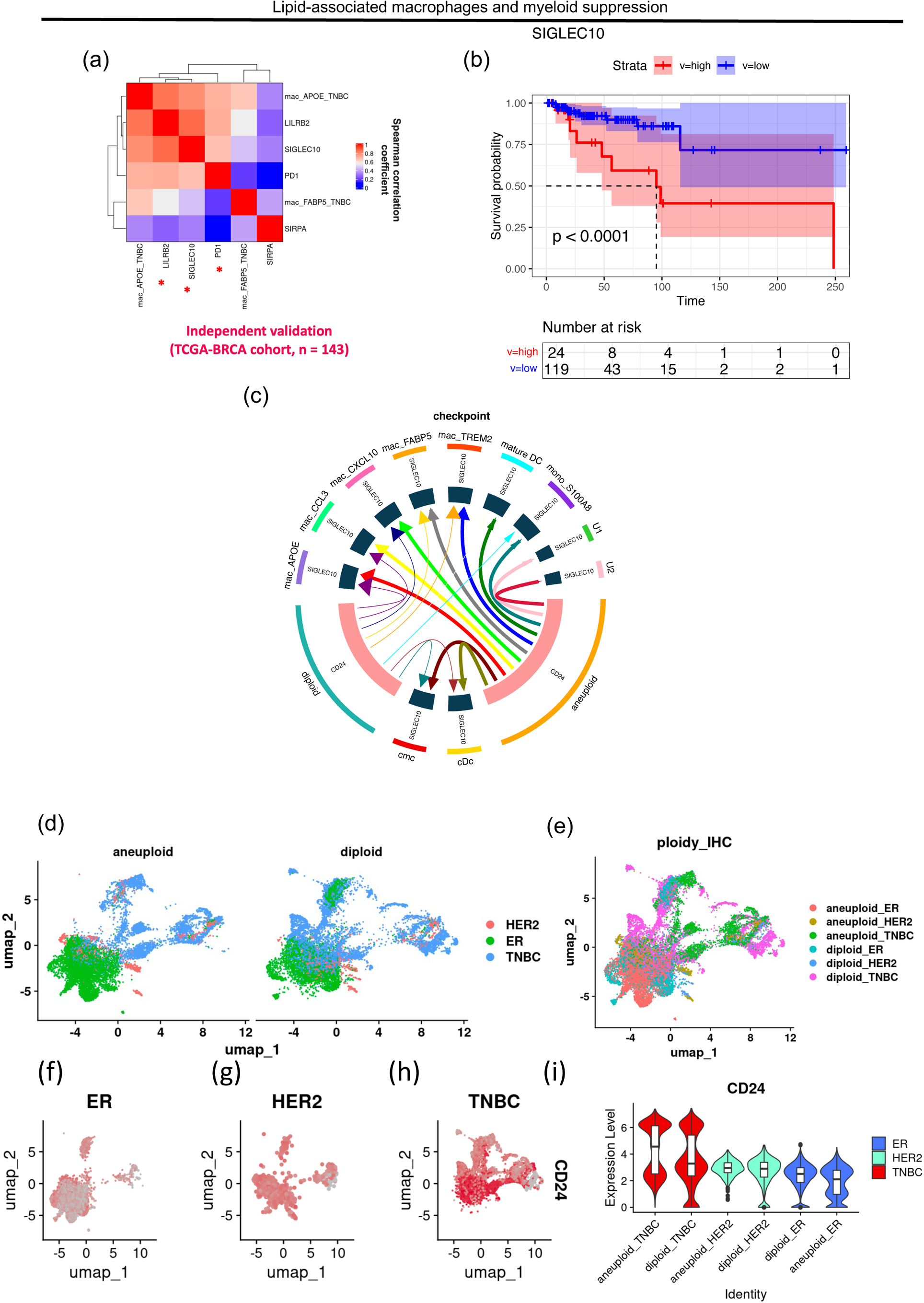
mac_APOE is associated with myeloid suppression via SIGLEC10 in TNBC. (a) Heatmap showing the Spearman correlation between the enrichment score of selected myeloid immune checkpoints and the lipid macrophage signature. The Spearman correlation coefficient (R) ranges from 0 to 1 in this heatmap. Colors of the heatmap correspond to the magnitude of R (red: 0.6 < R < 1; white: R = 0.6; blue: 0 < R < 0.6). (b) Kaplan-Meier survival analysis of the myeloid immune-checkpoint *SIGLEC10*. The dashed lines indicate the median survival time for the high-risk and low-risk cohorts. The 95% confidence interval bands for the high-risk and low-risk cohorts are shaded in low-opacity red and low-opacity cyan, respectively. The rectangle box below the curves contains count of individuals who are at risk over the given period. (c) Chord diagram illustrating the potential receptor-ligand interactions between myeloid subtypes and aneuploid epithelial cells in TNBC. (d-e) UMAP showing the overall distribution of diploid and aneuploid epithelial cells across different clinical subtypes of breast cancer. The ploidy of epithelial cells were distinguished using *copykat* R package. (f-h) Feature plot showing the differential expression of *CD24* in epithelial cells across different clinical subtypes of breast cancer. (i) The associated violin plots shows the expression level of *CD24* in diploid and aneuploid epithelial cells across different clinical subtypes of breast cancer. As for the boxplot inscribed within each violin plot, its mid-line represents the median of the expression score while the whiskers denote 1.5 x IQR.

We further investigated the importance of the above-mentioned myeloid checkpoints in modulating myeloid-mediated immunity in TNBC. First, *CD24* and *B2M*, which both ligand *SIGLEC10* and *LILRB2* respectively, were expressed at significantly higher levels in TNBC compared to ER^+^ and HER2^+^ breast cancer cells (Figure 4f-4i). The higher *CD24* expression in TNBC-derived cells is concordant with the earlier finding by Barkal et al. (2019). Additionally, we found that aneuploid epithelial cells express higher levels of *CD24* compared to their diploid counterparts (Figure 4i), implying that the genomic instability driven by aneuploidy may contribute to upregulated *CD24* expression in TNBC. Indeed, both *CD24* and *B2M* trigger "don’t eat me" signal in myeloid cells (Liu et al., 2023). In contrast, we found that PD-L1 (*CD274*), the ligand of the T-cell checkpoint PD1, is minimally expressed by the epithelial cells of all different subtypes (Supplementary Figure S4d), revealing its relatively minor effect in suppressing myeloid-mediated immunity in breast cancer including TNBC.

We further confirmed whether the *CD24/SIGLEC10* axis or the *LILRB2/B2M* axis suppresses myeloid-mediated immunity. Our cell-cell communication analysis using the *iTalk* R package (Wang et al., 2019; Supplementary Figure S4b) found that various TAM subtypes, including mac_APOE_TNBC and mac_FABP5_TNBC, were substantially regulated by aneuploid epithelial cells via the *CD24-SIGLEC10* axis (Figure 4c). Intriguingly, although *B2M* was upregulated in TNBC compared with other breast cancers, we did not find significant enrichment of the *LILRB2-B2M* axis between myeloid subtypes and aneuploid epithelial cells (Figure 4c). This suggests that the *LILRB2*-driven poor survival in basal-like breast cancer patients may be independent of the suppression of myeloid cells by tumor cells via the *LILRB2/B2M* axis. Altogether, the *CD24/SIGLEC10* axis is the major myeloid immune-checkpoint signal in TNBC.

### Immune checkpoint therapy reprograms lipid TAM to anti-tumour FOLR2^+^ TAM

We finally sought how to target lipid-associated TAMs. We analyzed the combinatorial treatment of atezolizumab (anti-PD-L1 antibody) and paclitaxel used in the IMpassion130 study (Schmid et al., 2020). scRNA seq data of TNBC samples from Zhang et al. (2021) dissected the myeloid transcriptomic landscape of treatment responders and non-responders (Figure 5a and Supplementary Figure S5b-S5d). The responder cohort reduced mac_APOE signature with the combined treatment (Figure 5b). In contrast, paclitaxel monotherapy could not reduce the mac_APOE signature (Figure 5c). This suggests that atezolizumab and paclitaxel combo could target the mac_APOE signature. The responder cohort reduced the inflammatory signature in mac_APOE (Figure 5d and 5e) with the combinatorial treatment. Intriguingly, responder cohorts possessed higher ratio of mac_APOE than non-responders, suggesting this population as a therapeutic target (Figure 5f and 5g).

**Figure 5.**
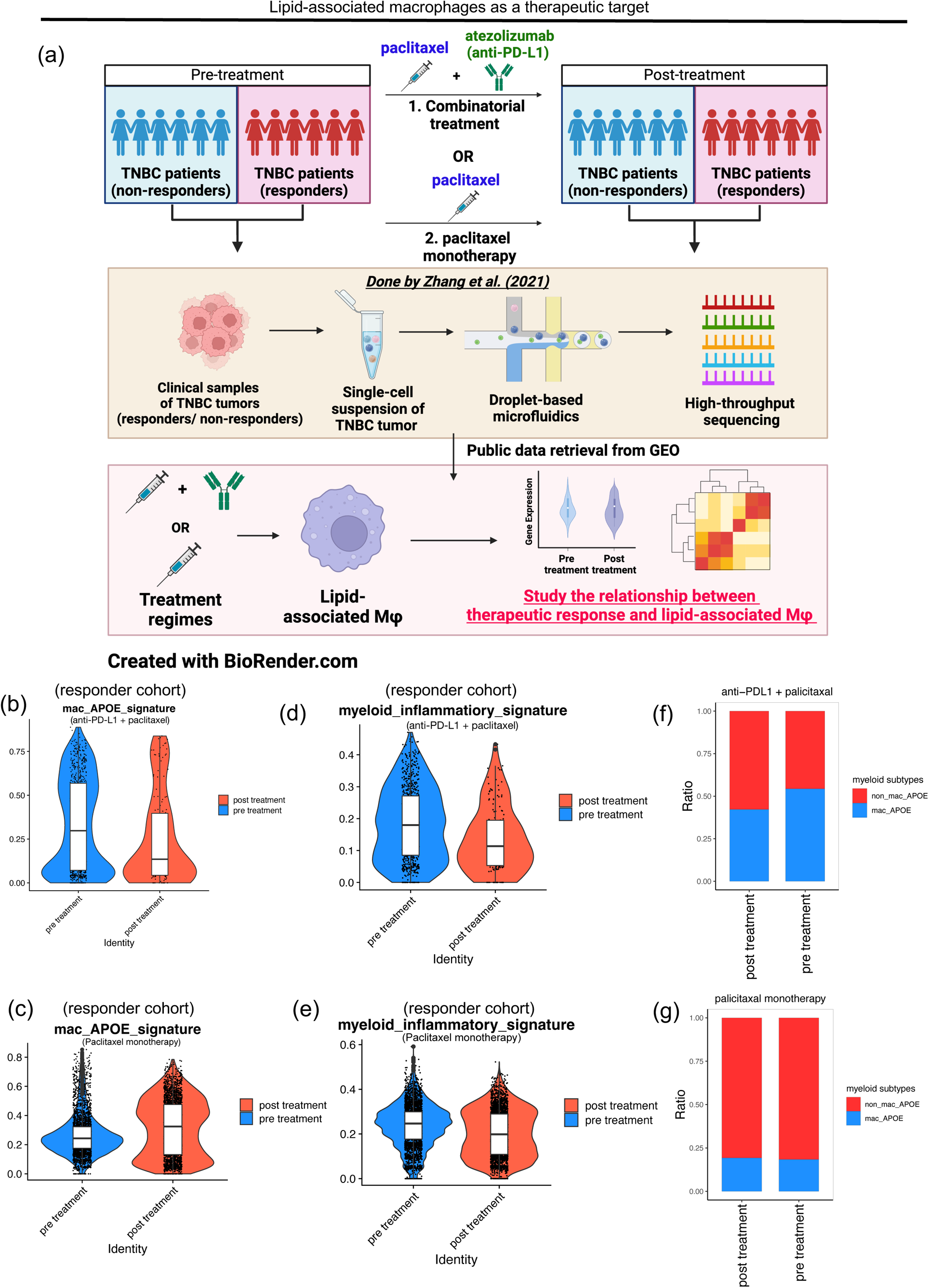
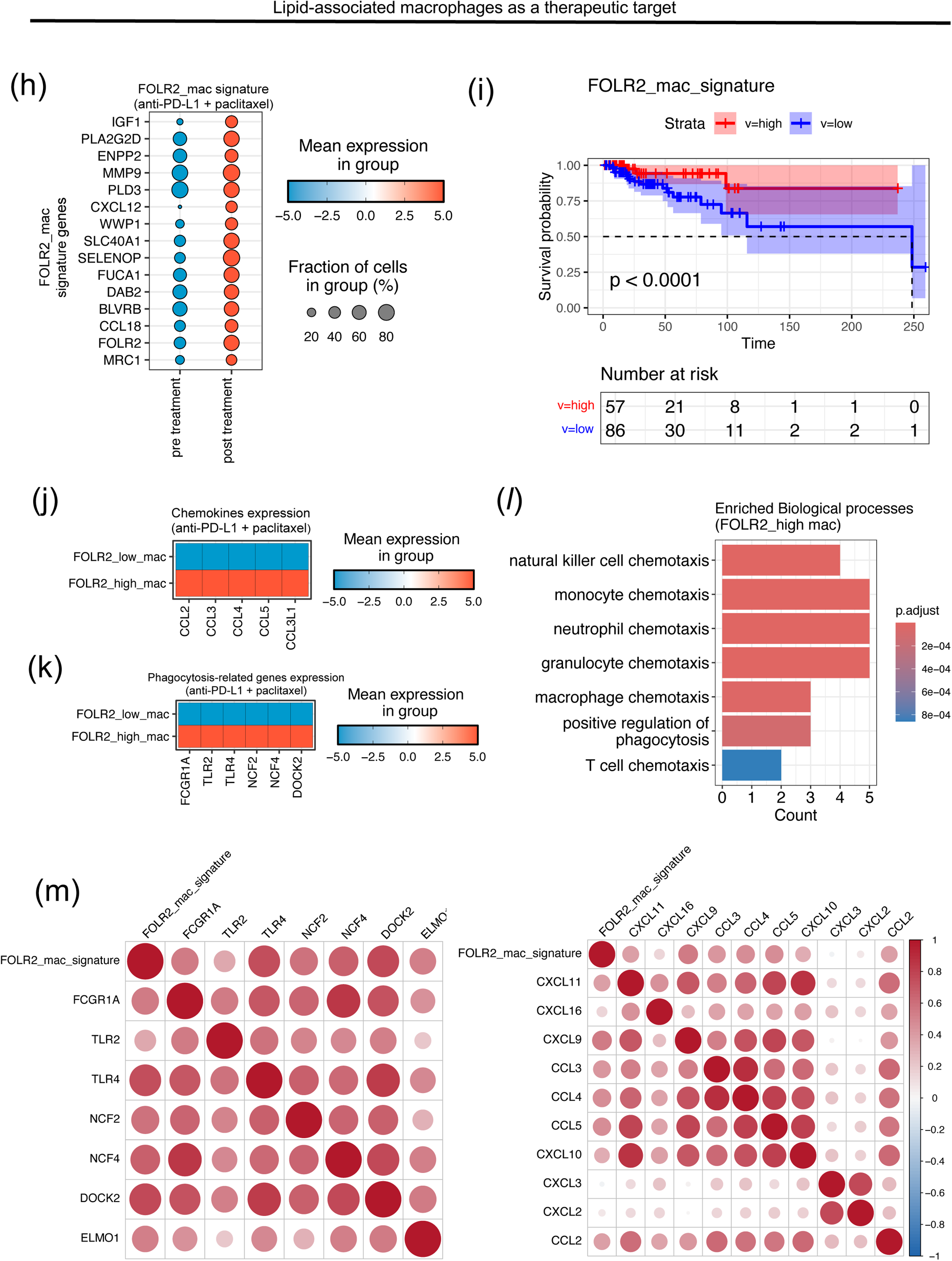
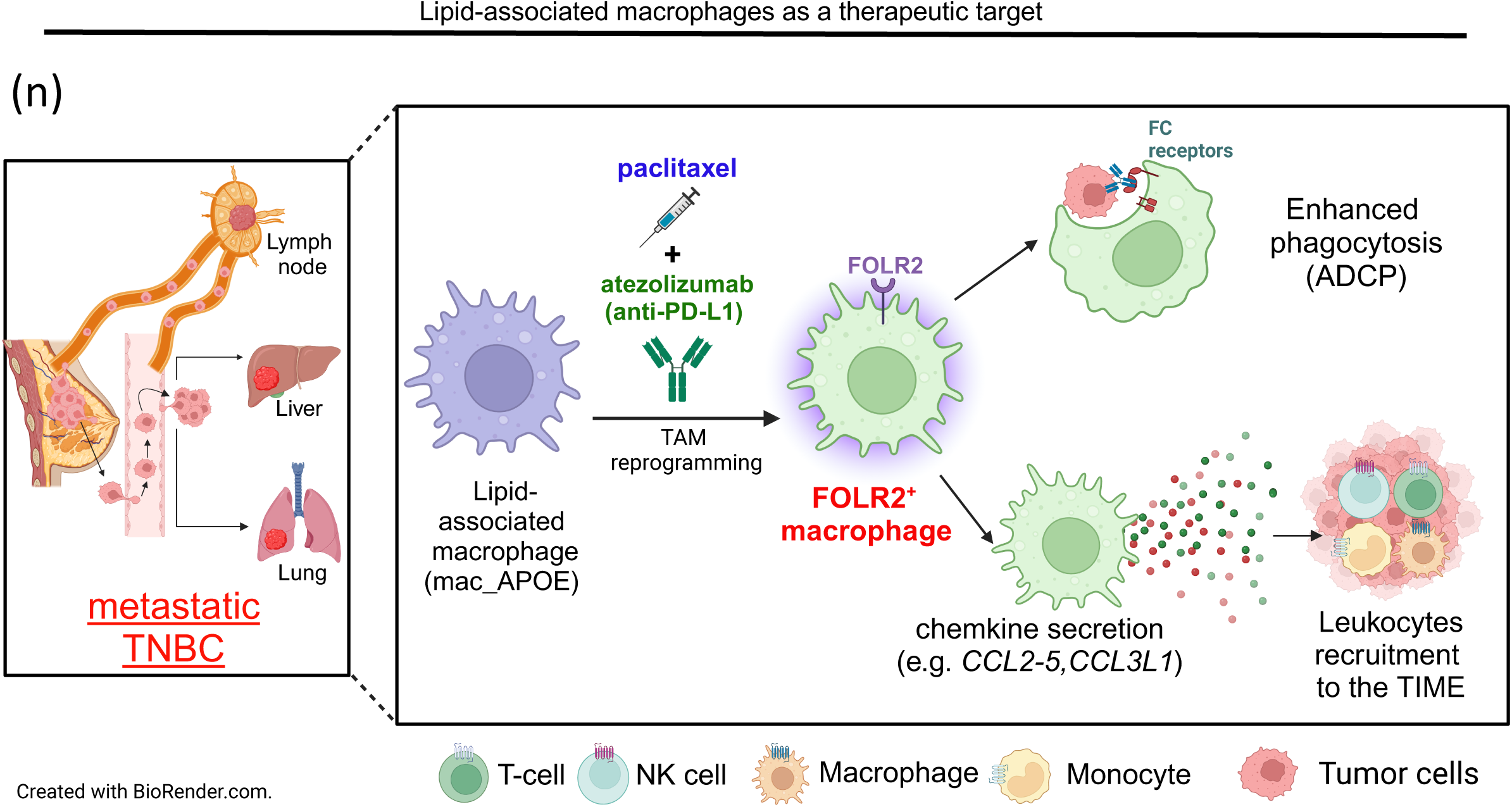
mac_APOE is a potential therapeutic target of combinatorial treatment using paclitaxel and atezolizumab. **(a)** Schematic diagram illustrating the overall experimental design for assessing the the therapeutic effect of the combinatorial therapy on the lipid-associated macrophages population. Created with BioRender.com **(b-c)** Violin Plots showing the expression of mac_APOE signature before and after receiving (b) combinatorial treatment and (c) paclitaxel monotherapy. The mid-line of each associated boxplot represents the median of the expression score while the whiskers denote 1.5 x IQR. **(d-e)** Stacked bar chart showing the relative proportion of mac_APOE and other types of TAMs (non_mac_APOE) before and after the (b) combinatorial treatment and (c) paclitaxel monotherapy. **(f-g)** Violin plots showing the expression of myeloid inflammatory signature associated with mac_APOE before and after receiving (b) combinatorial treatment and (c) paclitaxel monotherapy. The mid-line of each associated boxplot represents the median of the expression score while the whiskers denote 1.5 x IQR. **(h)** Bubble plot illustrating the upregulation of FOLR2^+^ macrophage gene signature in the mac_APOE cluster after combinatorial treatments **(i)** Kaplan-Meier survival analysis of the gene signature of FOLR2^+^ macrophages. The dashed lines indicate the median survival time for the high-risk and low-risk cohorts. The 95% confidence interval bands for the high-risk and low-risk cohorts are shaded in low-opacity red and low-opacity blue, respectively. The rectangle box below the curves contains count of individuals who are at risk over the given period. **(j-k)** Heatmaps showing the mean expression of chemokine genes and that of phagocytosis-related genes in the cluster of treatment-experienced mac_APOE, which consist of FOLR2_high and FOLR2_low macrophage after combinatorial treatment. **(l)** Barplot showing the enriched biological processes in the FOLR2_high macrophages after receiving the combinatorial treatment **(m)** Bubble plots illustrating the spearman correlation between the expression of FOLR2_mac gene signature, chemokine genes and phagocytosis-related genes. The correlation analysis was conducted using the bulk-RNA seq data of the basal-like breast cancer patients in the TCGA-BRCA cohort. **(n)** Schematic diagram illustrating the possible mechanism of how the combinatorial therapy involving atezolizumab and paclitaxel remodels the pro-tumorigenic mac_APOE in metastatic TNBC. Created with BioRender.com

We further dissected the transcriptomic landscape of the combinatorial-treatment-experienced mac_APOE clusters in the responder cohort. Intriguingly, we discovered two major sub-clusters of TAMs, namely FOLR2_high and FOLR2_low macrophages, where the FOLR2_high macrophages were characterized by the upregulation of *IGF1, PLA2G2D, ENPP2, MMP9, PLD3, CXCL12, WWP1, SLC40A1, SELENOP, FUCA1, DAB2, BLVRB, CCL18, FOLR2* and *MRC1* (Figure 5h and Supplementary Figure S5e), as reported by Nalio Ramos et al. (2022). Note that the upregulation of FOLR2_macrophage gene signature was not observed in most of the the non-responders (Supplementary Figure S5h), indicating that the FOLR2_high macrophage skewing might be the hallmark of the treatment response. Although Li et al. (2024) previously reported FOLR2_high macrophage as one type of oncofoetal cells in hepatocellular carcinoma (HCC), in breast cancer, FOLR2^+^ macrophages suggest better infiltration of immune cells, enhanced prognosis, and improved survival outcome in breast cancer (Nalio Ramos et al., 2022; Figure 5i). Single-cell transcriptomic analysis revealed that FOLR2_high macrophages elevated chemokines including *CCL2*, *CCL3*, *CCL4*, *CCL5* and *CCL3L1* (Figure 5j), which are key recruiters of leukocytes, such as T-cells, NK-cells and macrophages (Figure 5l). FOLR2_high macrophages highly expressed phagocytosis (*FCGR1A, TLR2, TLR4, NCF2, NCF4 and DOCK2*) (Figure 5k, 5*l*). Spearman correlation of the TCGA-BRCA cohort revealed a strong positive correlation between the mac_FOLR2 signature, chemokines, and the phagocytosis-related genes (Figure 5m). Altogether, the combinatorial treatment could reprogram the mac_APOE towards the FOLR2 macrophages to trigger functional immune responses against TNBC tumours (Figure 5n).

## Discussion

TNBC is well-known for its high aggressiveness and its association with a poor prognostic outcome. According to the earlier study by Pan et al. (2020), the immuno-suppressive milieu of TNBC is primarily driven by TAMs, which are responsible for the elevated production of immuno-suppressive cytokines, the reduced production of reactive oxygen species (ROS), and the increased infiltration of T-regs into the TIME. Here, we have also described a subtype of TAMs, known as lipid-associated macrophages, that demonstrate significant association with to the poor survival and immuno-suppressive phenotype in TNBC. We have uncovered a link between immunosuppression and the pro-inflammatory lipid-associated macrophages, shedding light on revealing the potential mechanisms by which the underlying detrimental association is caused. Our findings revealed that higher expression of lipid-associated macrophage signature corresponds to the T-cell exhaustion signatures, including increased expression of T-cell immune checkpoints, increased expression of transcription factors T-BET, TOX and EOMES, which are involved in driving T-cell exhaustion program. The close association between T-cell exhaustion and lipid-associated macrophages is further evidenced by the high probability of antigen presentation between lipid macrophages and CD8^+^ T-cells, as well as the correspondence of elevated expression of antigen processing machinery to higher expression of the lipid-associated macrophage signature.

Furthermore, we have identified positive correlation between elevated expression of lipid-associated macrophage signature and the expression of myeloid immune checkpoints like *SIGLEC10,* PD1 (*PDCD1*) and *LILRB2* and we have illustrated that *CD24/SIGLEC10* axis plays a relatively important role of myeloid immunosuppression in TNBC. Although the earlier study by Barkal et al. (2019) has illustrated the *in vivo* role of *CD24* in dampening phagocytic clearance of human breast cancer cell lines by TAMs using mouse model, the exact subtypes of TAMs regulated by *CD24/ SIGLEC10* derived from primary tumor have not been elucidated yet. Here, we have identified that not only lipid-associated macrophages are regulated by the *CD24/SIGLEC10* axis, but also other myeloid subtypes, such as mac_CCL3, mac_CXCL10, mac_TREM2, cDc, and cmc. However, other immune-checkpoints ligand, such as PDL1 and B2M, may play a less important role in suppressing myeloid-mediated immunity in TNBC. This illustrates the tremendous potential of *CD24/SIGLEC10* axis of being a promising target for immune checkpoint blockade for cancer immunotherapy. Altogether, this study has discovered that lipid-associated macrophages may play a crucial role in the immunosuppression TIME of TNBC, which could be the culprit of immune evasion and therefore the aggressiveness of TNBC. Nevertheless, the exact molecular mechanism of lipid-associated macrophage-dependent T-cell immunosuppression, as well as the manifestation of *CD24/SIGLEC10* axis, are still awaiting further studies.

Another significant finding from this work was the ability of the combinatorial therapy involving paclitaxel and atezolizumab to skew the pro-tumoral mac_APOE macrophages towards the phenotype of the pro-survival FOLR2_high macrophage, which were thought to be involved in the upregulation of genes related to chemotactic recruitment of immune cells, and those important for phagocytic activity. Such therapy-induced modification of macrophage phenotypes appears to be an essential mechanism by which the combinatorial therapy facilitates the orchestration anti-tumor immunity in TNBC. The above findings have briefly illustrated the complex interplay between drug response and the immune cell composition within the TIME, which may shed the light on developing more effective pharmacological strategies for better clinical outcome of TNBC patients. Nevertheless, additional research into the precise molecular processes that drive the modification of TAM’s phenotype might inform the design of more targeted therapy approaches.

Regarding the limitation, owing to the relatively small number of patients in all three cohorts of sc-RNA seq data used, the statistical power of the prediction of the molecular signatures and that of the therapeutic response might be limited, even though the correlations between molecular signatures were cross-validated using TCGA-BRCA bulk RNA-seq data. Additionally, since the locally advanced breast tumors from the TNBC patients receiving combinatorial treatment had been removed beforehand via surgery, only the metastasized tumor mass was available for single-cell transcriptomic analysis. The transcriptome of the metastasized TNBC may not truly reflect the TIME of the breast-localized TNBC, and hence, further analysis of how pharmaceuticals remodel the TIME of the breast-localized tumor would be necessary.

In addition, though our study has provided a thorough analysis of the transcriptomic landscape of the TIME of TNBC, with the focus on the mac_APOE and its skewness to FOLR2^+^ mac, further wet lab validation of our bioinformatic analysis will still be necessary. For instance, in terms of expression of chemokines and phagocytosis-related proteins, their protein level has still not been validated empirically. Lastly, this study mainly utilizes public sc-RNA seq data to compute the intra-cellular communication networks of tumor-associated cells. To further delineate the localization and the contact-dependent interactions between lipid-associated macrophages, tumor cells and other subtypes of immune and stromal cells, spatial transcriptomic analysis can be further conducted. Indeed, one category of TIME is known as infiltrated–excluded (I–E) TIME, which is characterized by the exclusion of cytotoxic lymphocytes from the core of the tumor, localization of cytotoxic lymphocytes at the periphery of tumor mass, and retention of cytotoxic lymphocytes by fibrotic nests (Binnewies et al., 2018). Tumor types with I-E TIME generally have limited anti-tumor immunogenicity. Hence, a comprehensive spatial analysis of TNBC could provide understanding of whether the immunosuppressive nature of TNBC is related to the existence of I– E TIME.

## Key Resources Table

**Key Resources Table:**
| Reagent or Resources | Source | Identifier |
| --- | --- | --- |
| **Deposited Data** |  |  |
| Processed sc-RNA seq data of 19 invasive ductal carcinoma patients | Wu et al. (2021) | GEO: [GSE176078](https://www.ncbi.nlm.nih.gov/geo/query/acc.cgi?acc=GSE176078) |
| Processed sc-RNA seq data of 8 TNBC patients | Pal et al. (2021) | GEO: [GSE161529](https://www.ncbi.nlm.nih.gov/geo/query/acc.cgi?acc=GSE161529) |
| Processed sc-RNA seq data of TNBC patients for assessing response to combinatorial therapy | Zhang et al. (2021) | GEO: [GSE169246](https://www.ncbi.nlm.nih.gov/geo/query/acc.cgi?acc=GSE169246) |
| **Bulk RNA-seq data of the TCGA-BRCA cohort** | UCSC genome browser database<br>Analytics Hub of Breast Cancer Now Tissue Bank, United Kingdom (clinical information of patients) | **Bulk RNA-seq data:** https://gdc-hub.s3.us-east-1.amazonaws.com/download/TCGA-BRCA.htseq_counts.tsv.gz<br>**Survival information:** https://gdc-hub.s3.us-east-1.amazonaws.com/download/TCGA-BRCA.mutect2_snv.tsv.gz<br>**Phenotypic information:** https://gdc-hub.s3.us-east-1.amazonaws.com/download/TCGA-BRCA.GDC_phenotype.tsv.gz<br>**Analytics Hub of Breast Cancer Now Tissue Bank, United Kingdom:** https://bcntb.bcc.qmul.ac.uk/analytics/cohort/tcga/ |
| **Software and algorithms:** |  |  |
| **RStudio CPU server** | Centre for Translational Stem Cell Biology (CTSCB), Hong Kong Science and Technology Park (HKSTP) | https://www.hkstemcell.hk/ |
| **RStudio (Version 4.3.2)** | R Core Team (2021) | https://www.r-project.org/ |
| **Seurat (Version 5.0.1)** | Satija et al. (2015) | https://satijalab.org/seurat/ |
| **Harmony (Version 1.2.0)** | Korsunsky et al. (2019) | https://github.com/immunogenomics/harmony |
| **COSG (Version 0.9.0)** | Dai et al. (2022) | https://github.com/genecell/COSGR |
| **iTalk (Version 0.1.0)** | Wang et al. (2019) | https://github.com/Coolgenome/iTALK |
| **CellChat (Version 2.1.2)** | Jin et al. (2021) | https://github.com/sqjin/CellChat |
| **scRNAtoolVis (Version 0.0.7)** | Zhang (2022) | https://github.com/junjunlab/scRNAtoolVis, https://junjunlab.github.io/scRNAtoolVis-manual/ |
| **Complexheatmap (Version 2.18.0)** | Gu et al. (2016) | https://github.com/jokergoo/ComplexHeatmap |
| **ClusterProfiler (Version 4.10.1)** | Yu et al. (2012) | https://github.com/YuLab-SMU/clusterProfiler |
| **org.Hs.eg.db (Version 3.18.0)** | Carlson (2019) | https://bioconductor.org/packages/release/data/annotation/html/org.Hs.eg.db.html |
| **survival (Version 3.5-8)** | Therneau et al. (2020) | https://cran.r-project.org/web/packages/survival/index.html |
| **survminer**<br>(Version 0.4.9) | Kassambara et al. (2021) | https://cran.r-project.org/web/packages/survminer/index.html |
| **copykat**<br>(Version 1.6.12) | Gao et al. (2021) | https://github.com/navinlabcode/copykat |
| **GSVA**<br>(Version 1.50.1) | Hänzelmann et al. (2013) | https://github.com/rcastelo/GSVA |
| **Others:** |  |  |
| **Online single-cell breast cancer atlas** | Wu et al. (2021) and Single Cell Portal, Broad Institute of MIT and Harvard. | [https://singlecell.broadinstitute.org/single_cell/study/SCP1039/a-single-cell-and-spatially-resolved-atlas-of-human-breast-cancers?genes=CD24&cluster=Myeloid%20cells&spatialGroups=-&annotation=subtype--group--study&subsample=all&tab=dotplot&scatterColor=.](https://singlecell.broadinstitute.org/single_cell/study/SCP1039/a-single-cell-and-spatially-resolved-atlas-of-human-breast-cancers?genes=CD24&cluster=Myeloid%20cells&spatialGroups=-&annotation=subtype--group--study&subsample=all&tab=dotplot&scatterColor=) |
| **BioRender** | BioRender | [BioRender.com](https://www.biorender.com) |

## Methods

### Sex as a biological variable

As for the sc-RNA seq datasets from Wu et al. (2021), Pal et al. (2021) and Zhang et al. (2021), all the patients from which the breast tumor samples were derived are of female sex. Hence, sex was NOT regarded as a source of biological variation in this study.

### Computational resources

All the computation analysis was conducted using RStudio CPU Server (Version 4.3.2 "Eye Holes", released on 31/10/2023) provided by the Centre for Translational Stem Cell Biology (CTSCB), Hong Kong Science and Technology Park (HKSTP). Terminal on MacOS was used to download the feature-barcode matrices of sc-RNA seq data.

### Selecting patients’ sc-RNA seq data for downstream analysis

As for the sc-RNA seq datasets of Wu et al. (2021), 19 female patients (CID3586, CID3838, CID3921, CID3941, CID3946, CID3948, CID4040, CID4067, CID4290A, CID44041, CID4461, CID4463, CID4465, CID4495, CID44971, CID44991, CID4515, CID45171 and CID4530N) were selected for downstream transcriptomic analysis. The average and median ages of the patients were 57.5 and 54, respectively. All the selected patient cohorts were treatment-naïve, such that any unwanted alteration of transcriptomic dynamics on the tumour samples by chemotherapeutic treatment was minimized. Additionally, all patients had invasive ductal carcinoma as the cancer type. As for the further details about the clinical information of patients, please refer to the Supplementary Table 1 (https://static-content.springer.com/esm/art%3A10.1038%2Fs41588-021-00911-1/MediaObjects/41588_2021_911_MOESM4_ESM.xlsx) given by Wu et al. (2021).

### Data Availability

The processed sc-RNA seq data used in this project were deposited in the Gene Expression Omnibus (GEO) of National Center for Biotechnology Information (NCBI) by Wu et al. (2021). The accession number for this dataset is GSE176078. As for the unprocessed sc-RNA seq data, they were deposited by Wu et al. (2021) in the European Genome-phenome Archive (EGA), and the corresponding accession number is EGAS00001005173 (https://ega-archive.org/studies/EGAS00001005173). As for the TCGA-BRCA bulk RNA seq data of 143 patients, they were retrieved from the University of California Santa Cruz (UCSC) genome browser database (https://gdc-hub.s3.us-east-1.amazonaws.com/download/TCGA-BRCA.htseq_counts.tsv.gz). Additionally, the survival and phenotypic information of the BRCA cohort was retrieved from the UCSC genome database (Survival information: https://gdc-hub.s3.us-east-1.amazonaws.com/download/TCGA-BRCA.mutect2_snv.tsv.gz; Phenotypic information: https://gdc-hub.s3.us-east-1.amazonaws.com/download/TCGA-BRCA.GDC_phenotype.tsv.gz) and Analytics Hub of Breast Cancer Now Tissue Bank, United Kingdom (https://bcntb.bcc.qmul.ac.uk/analytics/cohort/tcga/). The R code used for the analysis in this can be provided upon request.

### Code Availability

The code used for the analysis in this study has been deposited on GitHub and can be accessed with the link https://github.com/billychan509/CLChan_TNBC_analysis.

### Downstream processing of sc-RNA seq data

The *Seurat* R package (version 5.0.1) was used for downstream processing and analysis of sc-RNA seq data from 19 patients (Satija et al., 2015). Quality control of the sc-RNA seq data (Supplementary Figure S1b-S1d) was conducted by retaining only the cells with a mitochondrial percentage (mitoPercent) lower than 20%, number of features (nFeature_RNA) greater than 200, and a unique molecular identifier (UMI) count (nCount_RNA) greater than 250 (Supplementary Figure S1b). These criteria, employed by Wu et al. (2021), aim to filter out dying cells or cells with poor quality, both of which are characterized by mitochondrial DNA contamination. Next, the sc-RNA seq data in the Seurat object underwent "*LogNormalize*" normalization. This method normalizes the feature expression per cell with respect to the total feature expression and multiplies it by a scale factor of 10,000. Subsequently, the *FindVariableFeature* function from the *Seurat* R package was used to identify features that show a high degree of variation among cells. These features would be the focus of subsequent analyses on the variation in expression of biological features. The *ScaleData* function from the Seurat package was then used to perform a linear transformation of the dataset. Following this, the *RunPCA* function from the R package *Seurat* was applied to conduct Principal Component Analysis (PCA) on the scaled dataset. The *DimHeatMap* and *ElbowPlot* functions from the R package *Seurat* were used to generate dimensional reduction heatmaps and an elbow plot, respectively. These visualizations were used to determine which principal components (PCs) primarily contributed to the heterogeneity of the integrated datasets (Supplementary Figure S1e, S1g and S1h). In this analysis, the first 50 PCs were included in the downstream analysis.

### Integration of sc-RNA seq datasets and batch effect removal

During integration of sc-RNA seq data of 19 patients, R package *Harmony* (Version 1.2.0) was used to remove batch effect so that any technical variations, such as the experimental conditions during the preparation of sc-RNA library, that can mask actual biological variations between different (sub)types of cells can be minimized (Korsunsky et al., 2019). Hence, the chance that cells of the same biological types being scattered into different clusters on UMAP can be minimized.

### Identification of differentially expressed genes

The *FindAllMarkers* function from Seurat package (Satija et al., 2015) was employed to detect the differentially expressed features among various cell clusters, by leveraging the non-parametric Wilcoxon rank sum test. The annotations of cell (sub)clusters were based on the expression of the marker genes described above. Only genes with adjusted p-value (p-adjusted) lower than 0.05 were retained for further analysis.

Another R package, *COSG* (COSine similarity-based marker Gene identification), was further utilized to identify a list of characteristic marker genes expressed by the pre-categorized immune and stromal subclusters (Dai et al., 2022). The mathematical explanations for COSG are given below:

Let X be the N times M gene expression matrix obtained from *COSG*, in which N is equal to the total count of cells and M is the number of genes present in X. Then, vector *g_i_* is defined as the expression of *i^th^* gene in all the cells present in the matrix X’s *i^th^* column (equation 1), and *c_j_* denotes the expression level of *g_i_* in *j^th^* cell, where *j* belongs to the set of positive integers ranging from 1 to N. During marker gene identification, another variable*, λ_k_*, is defined as another artificial marker gene corresponding to Group *K* (*G_K_*), where *K* represents the number of cell clusters pre-categorized by other cell-annotation methods and belongs to the set of positive integers ranging from 1 to *K* (equation 2). If *c_j_* belongs to *G_k_*, the value of *λ_jk_* is set to 1. Conversely, if *c_j_* does not belong to *G_k_*, the value of *λ_jk_* is set to 0.

*COSG* calculates the angle between two vectors that depict the gene expression pattern within a cellular space of n dimension, where each dimension corresponds to one cell.

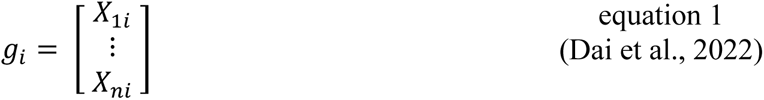

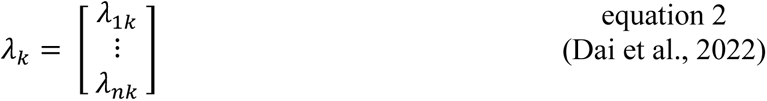

As for the vector that represents the expression of a specific gene, the number of bases corresponds to the total counts of cells detected, denoted as *n*. The extent of gene expression is illustrated by the coordinate that corresponds to each basis.

Hence, the value of *cos*(*θ*) is equivalent to the cosine similarity between two genes, where *θ* denotes the angle between the *λ_k_* and *g_i_*. The cosine similarity between *λ_k_* and *g* is evaluated by equation 3:

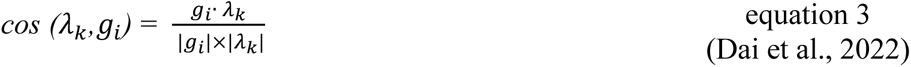

As for any values of *g_i_*, they would be selected as the marker genes for *Gk* if the value of *θ* attains a minimal value.

### Calculation of expression scores of selected gene signatures

The R package *Ucell* (Andreatta & Carmona, 2021) was used to quantify the expression of selected gene signatures. *Ucell* first obtained the relative ranks of genes in each of the cells, followed by calculation of Mann-Whitney U statistic (*U_j_*) (equation 4). The *UCell* score of selected gene sets for every cell *j* (*U*′*_j_*) can therefore be calculate by Equation 2 below:

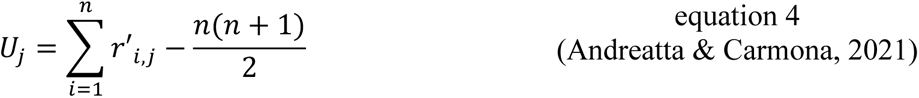

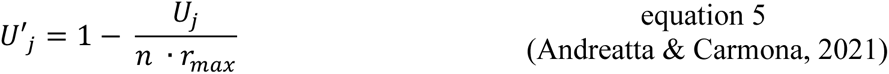

where *n* represents the number of genes present in a particular gene signature; *r’_i,j_* represents the relative rank of gene *i* in cell *j*; *r_max_* represents the threshold value determining the maximum rank of genes allowed in a cell. The default value of r_max_ is set to 1500 because during quality control processes, this threshold is commonly considered as the minimum requirement for the counts of genes detected in the dataset. As zero transcript counts are common in cell-gene matrix of sc-RNA seq because of the inability in capturing all transcripts in the original sample, which may lead to creation of a long chunk of lowest-ranked genes. To circumvent this problem, if *r_i,j_* > *r_max_*., *r_i,j_* will become *r_max_* + 1 such that all the lowest-ranked genes will be grouped up and provided a single rank value out of the range of valid ranks, which minimizes the computational demand on handling huge number of genes with lowest ranking.

### Identifying receptor-ligands pairs between selected genes clusters

Two R packages, namely *iTalk* (version 0.1.0) (Wang et al., 2019) and *CellChat* (version 2.1.2) (Jin et al., 2021), were utilized to identify potential cell-cell communication patterns in single-cell transcriptomic datasets.

As for the *iTALK* package, it was used to identify possible cell-cell communication network by mapping intercellular receptor-ligand pairs between selected cell clusters (Wang et al., 2019). The rawParse function was used to parse the normalized sc-RNA seq gene expression matrix and obtain the top 50% expressed receptor-ligand pairs. Then, LRPlot function was used to visualize the mean expression of inter-cellular receptor-ligand pairs by circos plot. The inter-cellular receptor-ligand pairs identified can be generally categorized into immune checkpoints, growth factors, cytokines, and other molecules.

As for the *CellChat* package (Jin et al., 2021), the CellChatDB.human, which is the built-in repository containing the receptor-ligands pairs expressed by human cells, was used as the database for inference of potential intercellular communication patterns in single-cell transcriptomic data. After discovering genes and receptor-ligand interactions that were overexpressed, the data of gene expression were projected, by diffusion process, onto an empirically validated network of protein-protein interaction (PPI) to lessen the impact of drop-out events present in the single-cell transcriptomic data. After that, the probability of communication among the interacting cell clusters were calculated by the built-in computeCommunProb function, where Tukey’s trimean was used to calculate the mean expression of genes in each group of cells. As for each group of cells, the minimum cell count needed for inter-cellular communication was set to 10.

### Identifying the gene signature of lipid-associated macrophages

The R package *COSG* (version 0.9.0) (Dai et al., 2022) was used to identify the top 50 enriched genes in the mac_APOE_TNBC and mac_FABP5_TNBC clusters. The R functions intersect was used to identify the overlapping gene sets between the lipid-associated macrophages from two independent sc-RNA seq datasets (Pal et al., 2021; Wu et al., 2021). Theses overlapping genes were then denoted as the marker genes for lipid-associated macrophages. Venn diagrams, visually depicting the commonalities and differences in expressed genes between subpopulations of cells, were generated using the R function venn.diagram provided by the R package *VennDiagram* (Version 1.7.3) (Chen & Boutros, 2011).

### Independent validation of molecular signatures of lipid-associated macrophages using other publicly available sc-RNA seq data

The identified molecular signatures of lipid-associated macrophages were further validated using another set of public sc-RNA seq data of TNBC published by Pal et al. (2021), which can be retrieved from NCBI using the accession number GSE161529. Eight TNBC patients, including Patient 0090, Patient 0126, Patient 0135, Patient 0106, Patient 0114, Patient 4031, Patient 0131, Patient 0554, and Patient 0177, were included in this validation cohort. The mean and median age of these eight patients were 55.25 and 62.5 respectively. The genuine molecular signatures of lipid macrophages were identified by comparing the top 50 genes expressed by the lipid-associated macrophage cluster in Wu et al. (2021) and that in Pal et al. (2021) (Figure 1g). As for the further details about the clinical information of patients, please refer to the Table EV2 of Pal et al. (2021). (https://www.embopress.org/doi/suppl/10.15252/embj.2020107333/suppl_file/embj2020107333-sup-0004-tableev2.docx).

### Characterization of myeloid immunosuppression and inflammatory signature

The genes encoding myeloid immune checkpoints (*LILRB2*, *SIGLEC10*, PD1(*PDCD1*) and *SIRPA*) were obtained from Zhou et al. (2020) and Liu et al. (2023) while the myeloid inflammatory signatures (*CCL11, IFNG, CCL3, CCL4, IL1B, TNF, IL1R2, CD80, CD86, TLR2, HIF1A, CSF2, IL6, CCL2, CCL8, CCL5, CD68, IRF5* and *TLR4*) were obtained from Zhang et al. (2021).

### Characterization of CD8+ T-cell exhaustion phenotypes

The exhaustion phenotype of CD8^+^ T-cells are characterized based on the exhaustion markers The exhaustion phenotype of CD8^+^ T-cells is characterized based on the enrichment of exhaustion markers/ T-cell suppression markers including T-cell immune checkpoints *(CTLA4, PDCD1, TIGIT, HAVCR2, LAYN*, and *LAG3*), as well as the expression of effector molecules (*IL2, TNF,* and *IFNG*). Additionally, transcription factors that drive T-cell exhaustion (*T-BET, TOX,* and *EOMES*) were obtained from Franco et al. (2020) and Belk et al. (2022) and were used to assess their correlation with the mac_APOE_TNBC signature.

### Clinical classification of patient samples by expression hormone receptors

The clinical subtypes of patient samples were determined via immunohistochemistry by Wu et al. (2021). The relevant experimental procedures were described by Wu et al. (2021). The clinical subtypes of each patient were obtained from the Supplementary Table 1 of Wu et al. (2021). (https://static-content.springer.com/esm/art%3A10.1038%2Fs41588-021-00911-1/MediaObjects/41588_2021_911_MOESM4_ESM.xlsx)

### Visualization of differentially expressed features

The functions from the R package *scRNAtoolVis* (version 0.0.7) (Zhang, 2022), including AverageHeatMap and jjdotplot, were used to visualize the Z-scores and mean expression levels of differentially expressed genes, respectively. FeatureCornerAxes function from *scRNAtoolVis* was also used to visualize the expression of selected genes on UMAP. The FeaturePlot function (Satija et al., 2015) from *Seurat* package was used to illustrate the subpopulations of cells on UMAP that possess high expression of selected genes., *VlnPlot* function from *Seurat* package (Satija et al., 2015) was used to create violin plots that depict the distribution of expression levels of selected genes /gene signatures. The *Complexheatmap* package (version 2.18.0) (Gu et al., 2016) was used to generate heatmaps that show the correlations between the selected gene signatures.

### Gene ontology (GO) analysis

GO analysis of the selected cell clusters were conducted using the R package *ClusterProfiler* (version 4.10.1) (Yu et al., 2012). Human genes were annotated in a genome-wide manner using the R package *org.Hs.eg.db* (version 3.18.0) (Carlson, 2019).

### Copy Number Variations (CNVs) analysis

The CNVs analysis of EPCAM^+^ epithelial cells were conducted using R package *copykat* (version 1.6.12) as described by Gao et al. (2021). The aneuploid EPCAM^+^ epithelial cells are distinguished from the diploid counterparts.

### Assessing the therapeutic response of TNBC patients to combinatorial therapy

The single-cell RNA sequence data of TNBC tumors before and after combinatorial treatment/paclitaxel monotherapy had been deposited by Zhang et al. (2021) and were obtained from GEO for our study using the accession number GSE169246. As for the combinatorial therapy, four responders (P007, P010, P012 and P019) and five non-responders (P002, P004, P005, P016, and P017) were chosen for assessing the therapeutic response. As for the paclitaxel monotherapy, three responders (P022, P020 and P013) and four non-responders (P025, P018, P023 and P003) were chosen for further analysis. To check whether the selected treatment regimens could target the pro-tumor mac_APOE, the relative expression scores of mac_APOE signature before and after treatments were compared. Additionally, the expression of myeloid inflammatory signature was assessed before and after the treatments. We also checked whether the mac_APOE of treatment-experienced patients would be biased towards the phenotype of FOLR2^+^ macrophage, which was characterized by the expression of *MRC1, FOLR2, CCL18, BLVRB, DAB2, FUCA1, SELENOP, SLC40A1, WWP1, CXCL12*, *PLD3*, *MMP9, ENPP2, PLA2G2D* and *IGF1*, as reported by Nalio Ramos et al. (2022). The expression of chemokines (*CCL2, CCL3, CCL4, CCL5* and *CCL3L1*) and that of phagocytosis-related genes (*FCGR1A, TLR2, TLR4, NCF2, NCF4* and *DOCK2*) in the FOLR2_high macrophage cluster were studied.

### Survival analysis using publicly available bulk RNA seq data of TCGA-BRCA cohort

Using the R packages *survival* (version 3.5-8) (Therneau, 2020) and *survminer* (version 0.4.9) (Kassambara et al., 2021), Kaplan-Meier survival analysis was conducted to assess the potential impact of immune cell and stromal cell populations on patient survival in the TCGA-BRCA cohort. Only patients exhibiting the basal phenotype, which resembles the TNBC phenotype characterized by the absence of ER, HER2, and PR expression, were retained for survival analysis (n = 143). Stringent patient selection criteria were applied, including the exclusion of patients without tumor tissue and those lacking survival information or surviving for fewer than 30 days. The R package *COSG* (Dai et al., 2022), as mentioned above, was utilized to identify differentially expressed marker genes among the selected immune and stromal cell subtypes. Regarding the use of *COSG* package, the parameter *expressed_pct* was set at 0.5 to ensure that the candidate marker genes are substantially expressed within the subclusters whilst *n_genes_user* was set as 50, allowing selection of the top-ranked gene having the most pronounced differential expression. Moreover, the penalty factor μ, was set as 1, which aims to penalize the gene expression in non-target subclusters, hence reducing the inclusion of housekeeping genes and enhancing the specificity of the identified marker genes for the analyzed subclusters. Such rigorous approach aimed to obtain biologically meaningful sets of marker genes expressed by the respective sub-clusters of tumor-associated cells, bettering subsequent survival analysis and interpretation.

The gene set enrichment scores of selected gene sets were calculated using the bulk RNA sequencing data of the TCGA-BRCA cohort, utilizing the *gsva* function from the *GSVA* R package (version 1.50.1) as described by Hänzelmann et al. (2013). The *surv_cutpoint* function, which utilizes maximally selected rank statistics, was used to identify the optimal cutpoint in the expression levels of the gene set that demonstrates the strongest association with patient survival. The *ggsurvplot* function was then employed to construct the survival plots, using the clinical information of the TCGA-BRCA cohort and the differentially expressed marker genes of the selected subtypes of tumor-associated immune cells as inputs for constructing the survival plot.

### Dichotomizing the basal-like breast cancer patients in TCGA-BRCA cohort based on the enrichment level of selected gene sets

The enrichment score of mac_APOE_TNBC gene set was dichotomized into groups known as “mac_APOE_TNBC High” if the enrichment score was greater than 0, and “mac_APOE_TNBC Low” if the enrichment score was lower than 0. These two categorical variables were then used to assess the differences in enrichment scores of selected gene sets.

### Statistics

The difference in median enrichment level of selected genes signatures (i.e. T-cell suppression signature, T-cell exhaustion-related transcription factors, antigen-processing machinery) between patients having higher expression of mac_APOE_TNBC signature and those having lower expression of mac_APOE_TNBC signature are assessed using Wilcoxon rank-sum test. Additionally, Kruskal-Wallis test was used to checking the difference in median of module scores of selected immune checkpoint ligands in epithelial cells across different clinical subtypes of breast cancer. The normality and the homogeneity of variance the enriched genes scores were assessed by Shapiro-Wilk Test and Levene’s Test, respectively.

Additionally, Spearman correlation analysis was performed using bulk RNA seq data from the TCGA-BRCA cohort to independently assess the correlation between the bimodally distributed gene set enrichment scores of selected gene signatures. As described above, the gene set enrichment scores were calculated using the *gsva* function from the *GSVA* R package (Hänzelmann et al., 2013). Each immune subtype was defined by the expression levels of the top 50 marker genes identified by *COSG*, as mentioned earlier (Dai et al., 2022).

Only associations with respective p-values lower than 0.05 will be considered statistically significant. Associations with a p-value less than 0.001 are annotated with the "***" symbol, indicating a highly significant relationship. Associations with a p-value between 0.001 and 0.05 (exclusive) are annotated with the "**" symbol, which represent a moderately significant relationship. Lastly, any association with p-value between 0.05 and 0.1 (exclusive) are annotated with the "*" symbol, indicating a weakly significant relationship. All the statistical analysis was conducted using R.

### Creation of figures/ illustrations and editing of figures

BioRender was used for generating figures or illustrations. Any figures or illustrations produced with BioRender can be identified with the phrase "Created with BioRender.com". Additionally, figures were edited using Microsoft PowerPoint and Notability.

## Author Contribution

This study was conceptualised by C.L.C, A.T, S.Z, J.W.H.W, Y.H, Y.M.C and R.S. The investigation was conducted by CLC. Bioinformatics analysis was conducted by C.L.C. The original draft was written by C.L.C, and was further reviewed and edited by Y.M.C and R.S. The study was supervised by Y.M.C, Y.H, J.W.H.W and R.S.

## Address correspondence to

Ryohichi SUGIMURA, L3-64, Laboratory Block, 21 Sassoon Road, Pokfulam, Hong Kong SAR, China; Phone: +852 3917 9269; Or to Yiming CHAO, L1-05, Laboratory Block, 21 Sassoon Rd, Pokfulam, Hong Kong SAR, China; Phone: +852 6341 5297;

