## Supplementary_Material for "Lipidic and senescent macrophages predict progression and response to combinatorial immunotherapy in triple-negative breast cancer"

Supplementary Figure S1

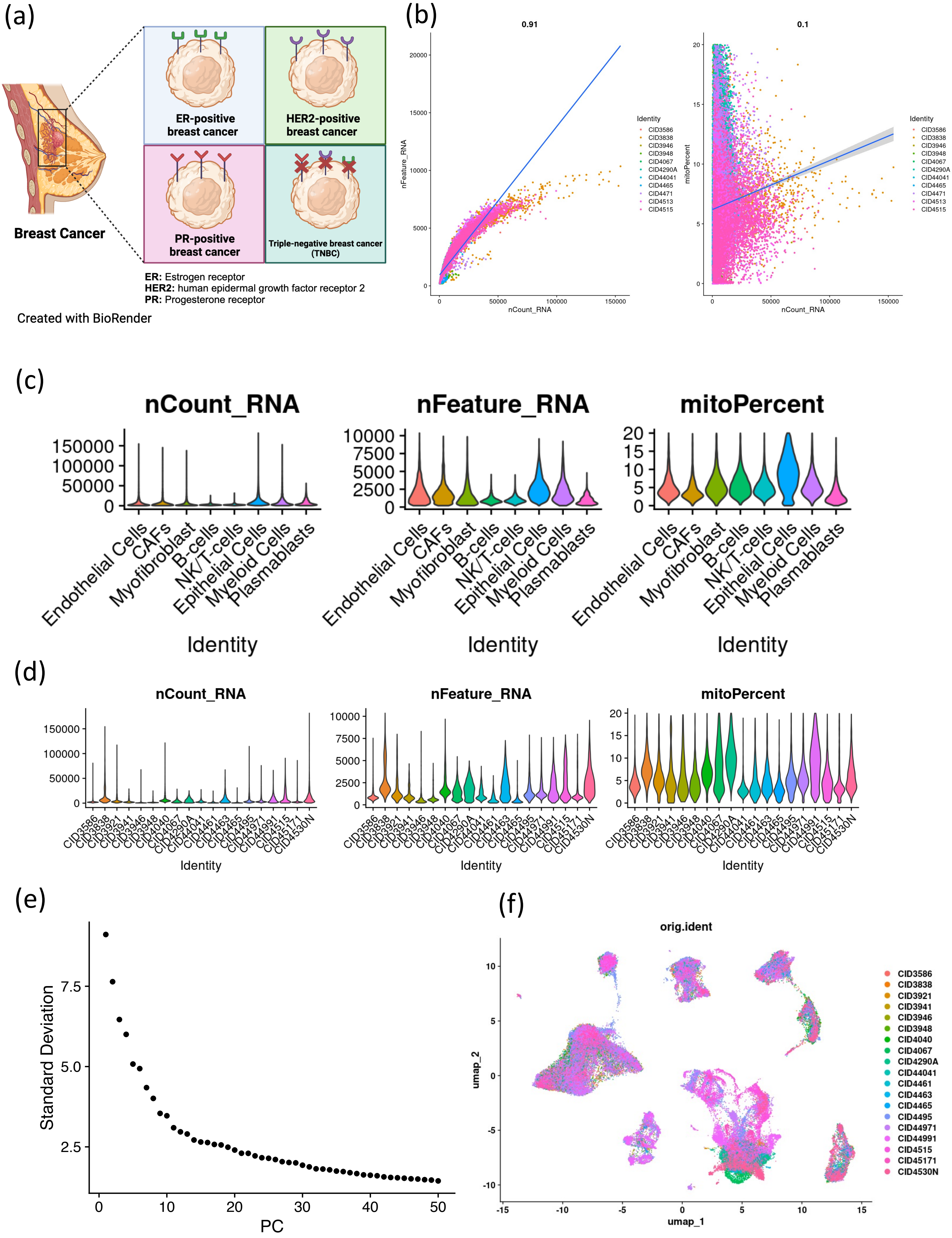

Supplementary Figure S1 (Contd)

(g)

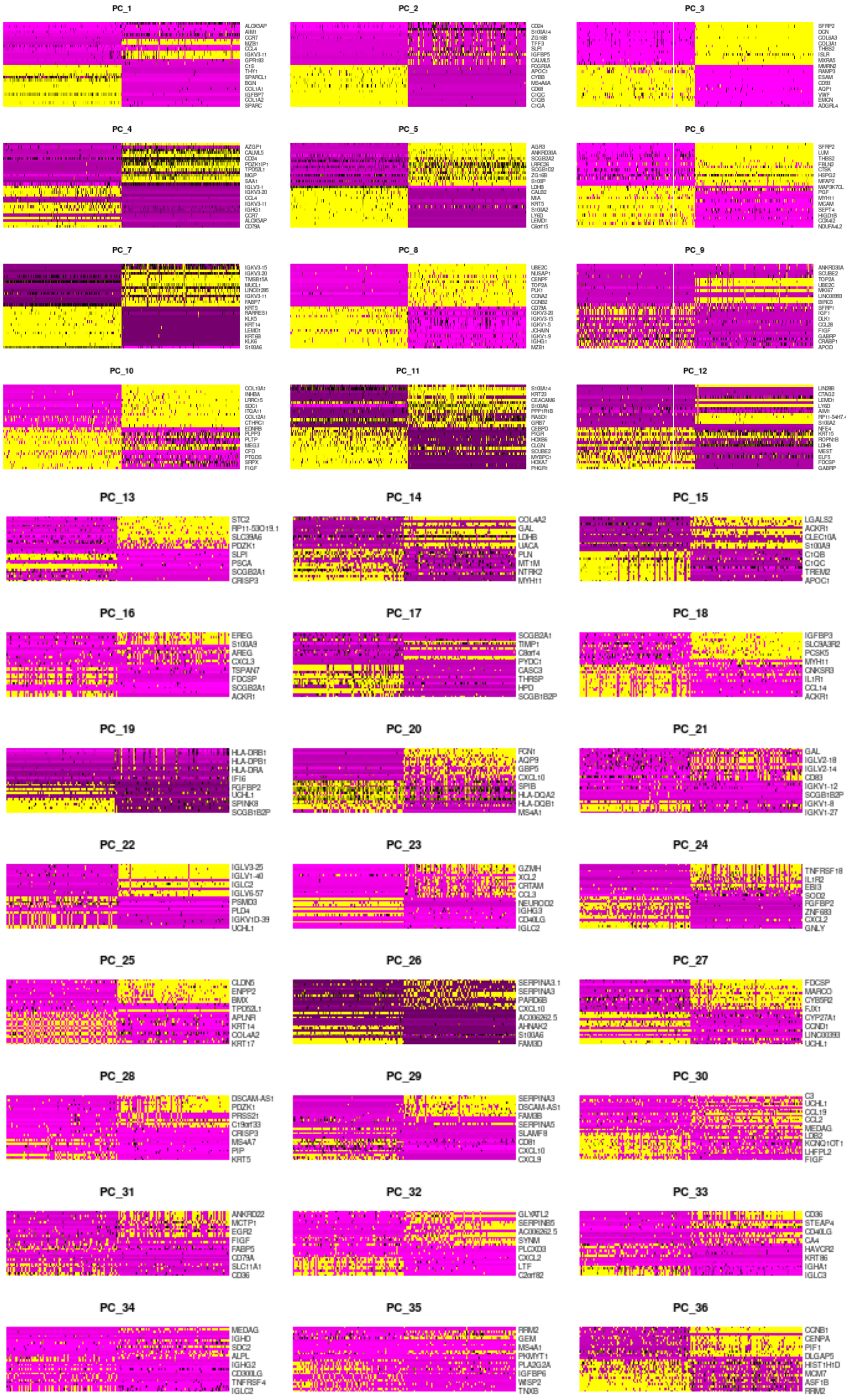

Supplementary Figure S1 (Contd)

(h)

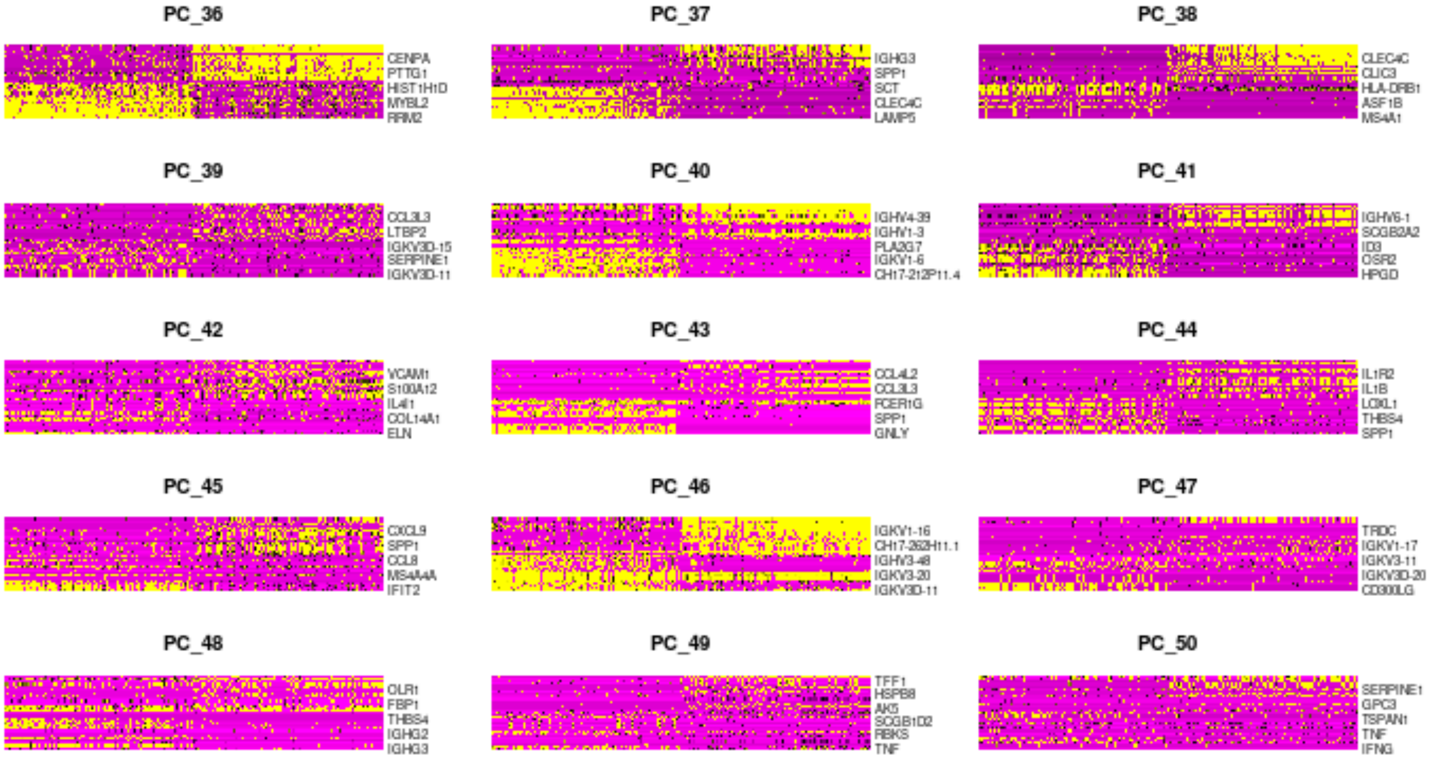


(j)

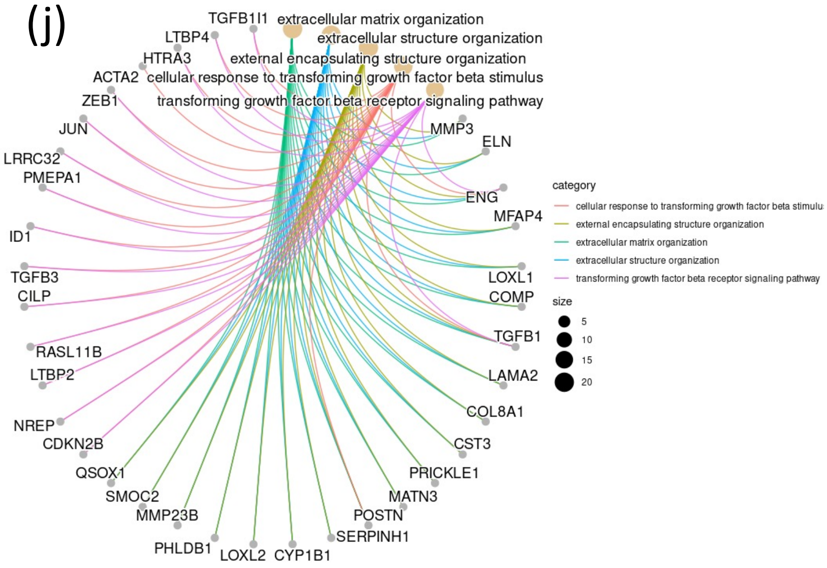


(k)

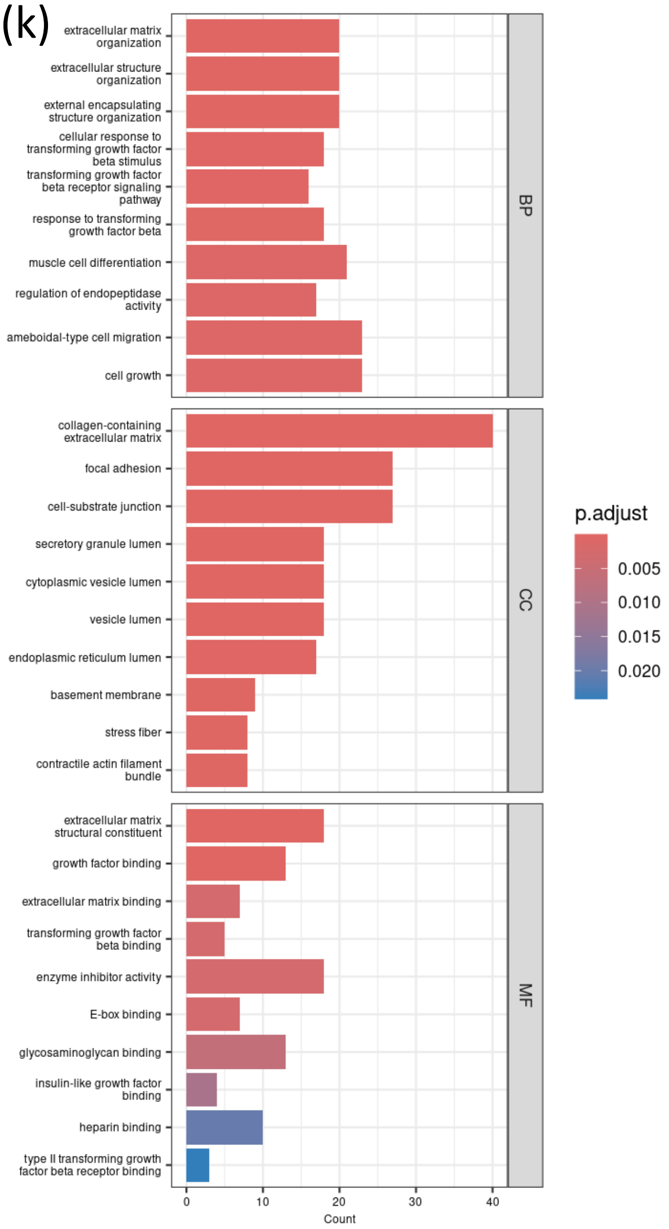


(*l*)

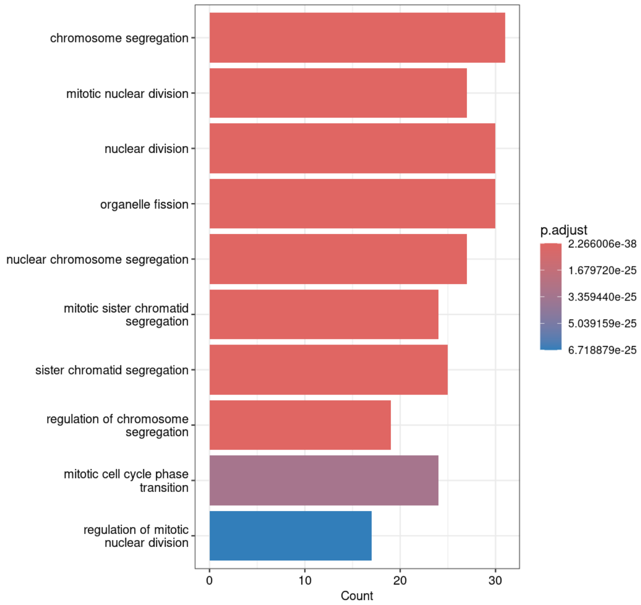

**Supplementary Figure S1**

(a) Cartoon illustrating the clinical classification of breast cancer based on hormone receptors (ER, HER2 and PR) expression. Breast cancer lacking the three types of receptors are known as triple-negative breast cancer (TNBC). Created with BioRender.com.

(b) Scatter plots depicting the correlation between (a) count of RNA molecules (nCount_RNA) and count of features (nFeature_RNA) and that between nCount_RNA and mitochondrial percentage

(c-d) Violin Plots showing the distribution of nCount_RNA, nFeature_RNA and mitoPercent across (a, b, c) different types of tumor-assoicated cells and (d, e, f) different patients of invasive ductal carcinoma after quality control. The scRNA seq dataset was adopted from Wu et al. (2021).

(e) Elbow Plot depicting the standard deviation of first 50 principal components in the scRNA seq data of 19 invasive ductal carcinoma patients from Wu et al. (2021)

(f) UMAP plot showing clusters of cells derived from 19 patients. Cell were categorized based on (a) patients' identity (orig.ident). The patient-derived tumors cells were found to be scattered in different clusters on UMAP, indicating the success in removal of batch effect.

(g-h) Heatmap illustrating the first 50 principal components in the scRNA seq dataset of Wu et al. (2021)

(j) Cnetplot showing the relationship between the enriched biological processes (BP) in matrix CAF and collagen CAFs and the relevant genes that contribute to the enrichment.

(k - *l*) Barplots showing the top 10 enriched BP cellular components (CC) and molecular functions in the matrix (k) CAF, collagen CAFs and (*l*) cycling myeloid cells (cmc)

Supplementary Figure S2

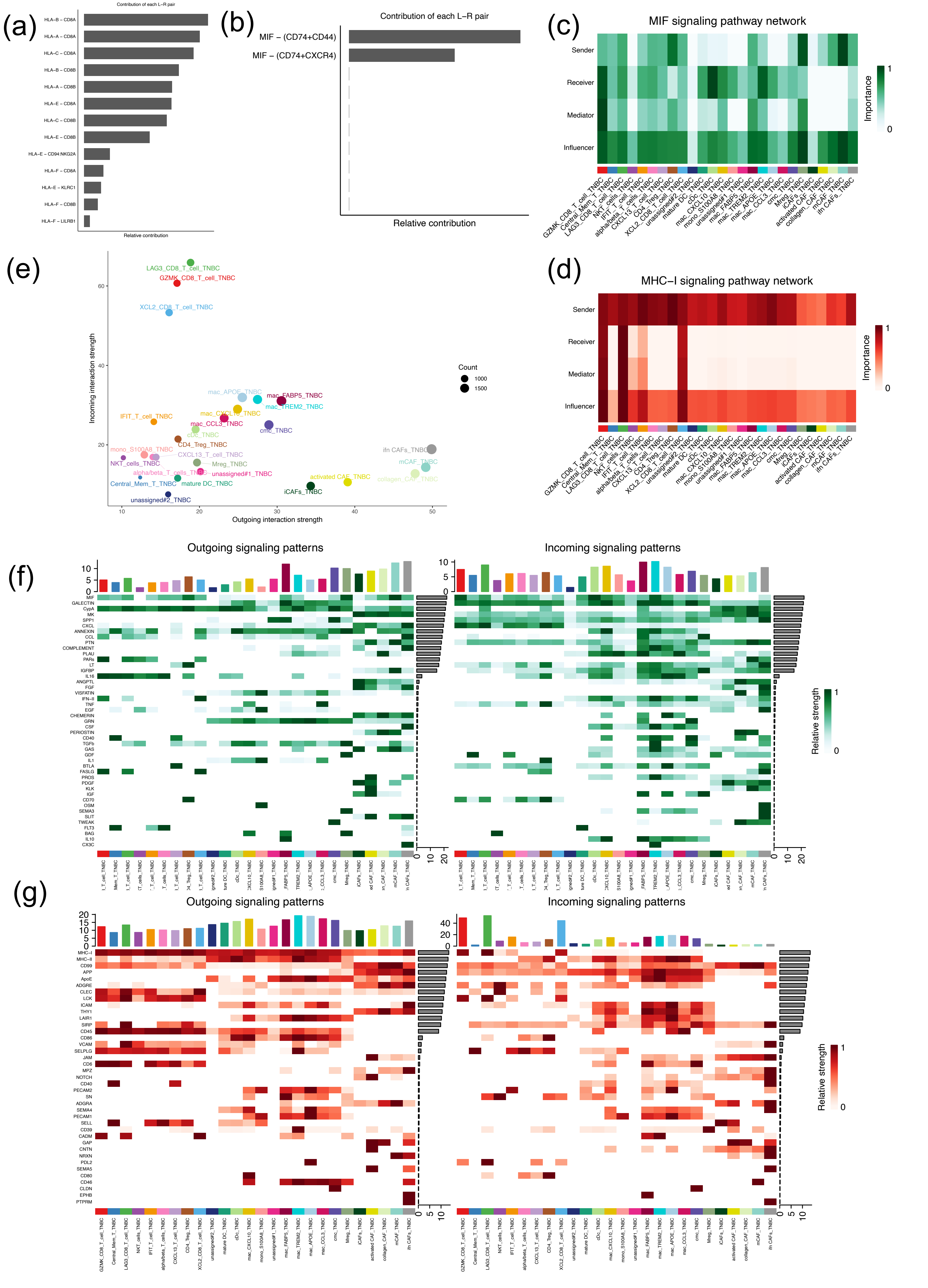

(h)

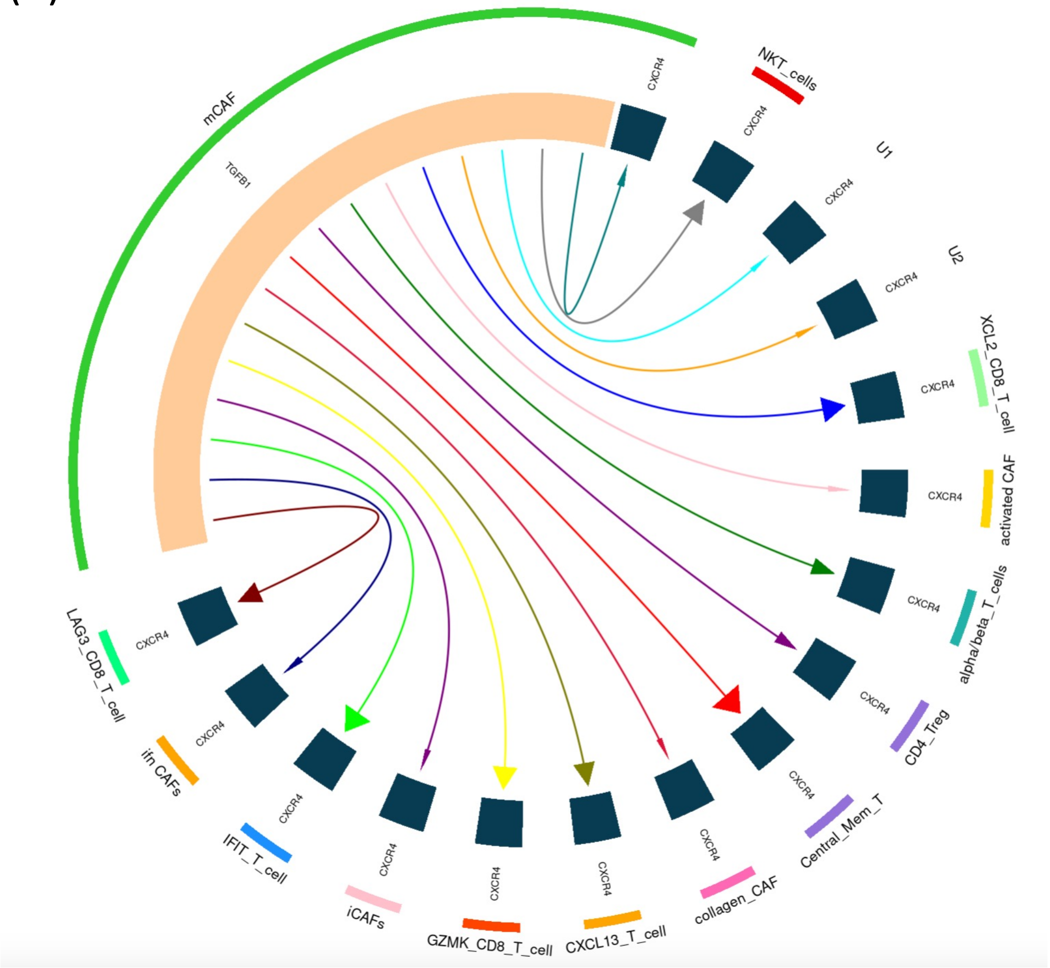


**Supplementary Figure S2**

(a-b) Barplots showing the relative importance of each ligand-receptor (L-R) pair in (a) MHC-I and (b) MIF signaling pathways.

(c-d) Heatmaps showing the senders, receivers, mediators and influencers in (c) MIF and MHC-I signaling pathway. The relative importance of each role in each type of tumor-associated cells is proportional to the color intensity.

(e) Bubble plot showing the relative incoming and outgoing interaction strength between different subtypes of tumor-associated cells. The size of each bubble is proportional to the count of the interactions in each subtype of cells.

(f-g) Heatmap showing the cell-cell communication patterns via (f) secreted signaling or (g) cell-cell contact between different types of tumor-associated cells.

(h) Receptor-ligand mapping between mCAF and tumor-associated T-cells via TGF-beta-CXCR4 axis. The receptor-ligand was constructed using *iTalk* R package

Supplementary Figure S3

(a)

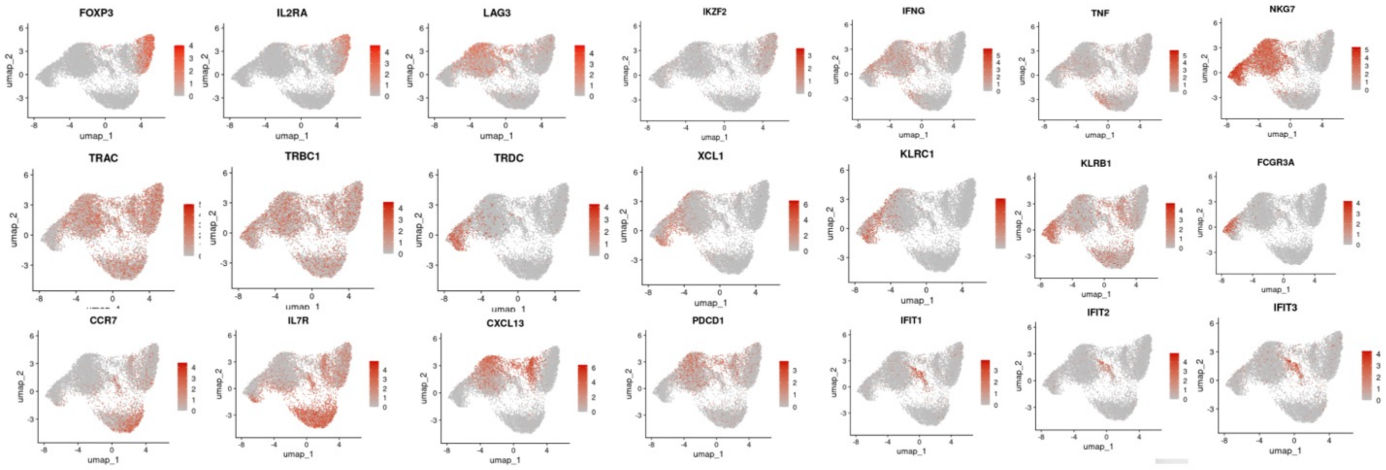


Marker genes for T-cells, NK-cells and NKT-cells subcluters

(b)

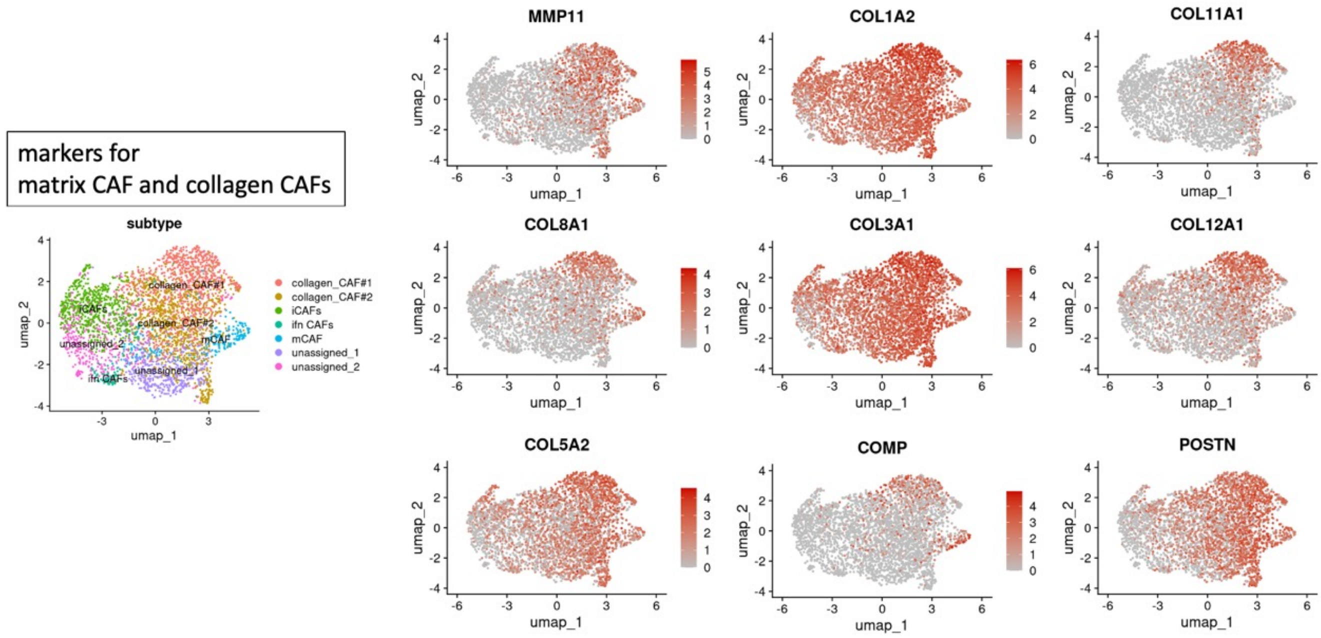


(c)

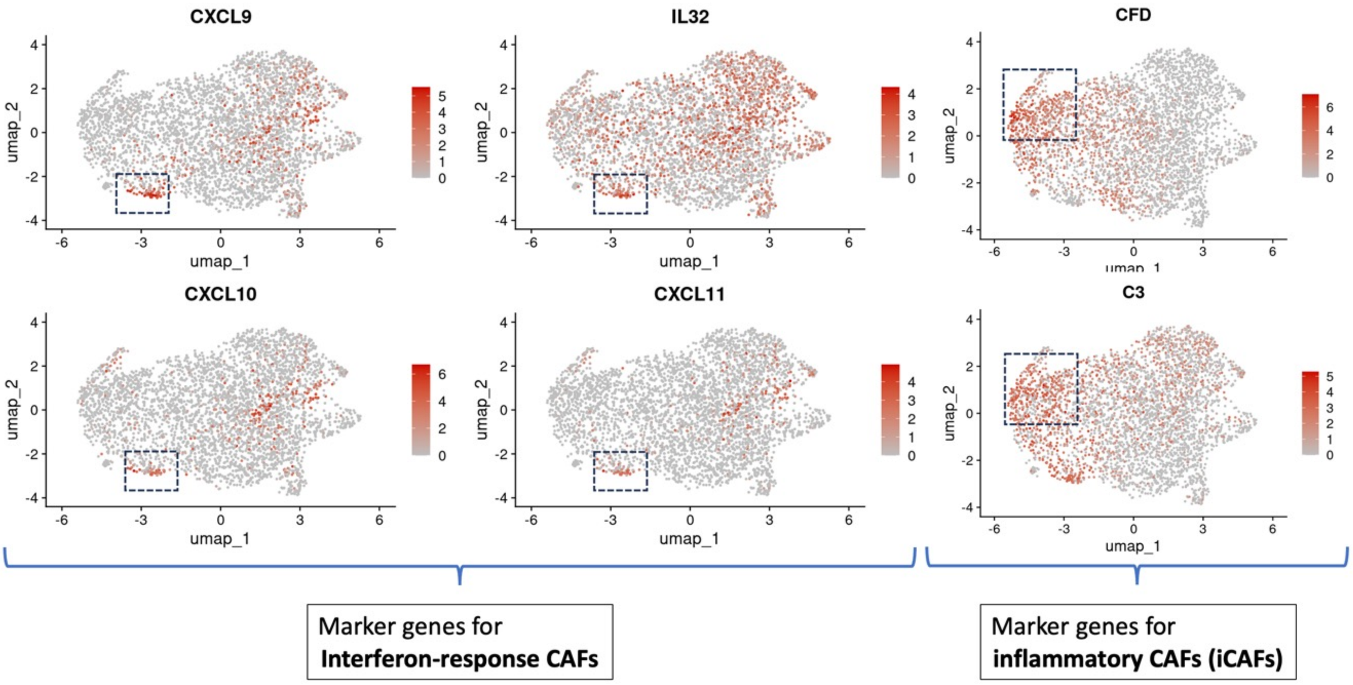

Supplementary Figure S3 (Contd.)

(d)

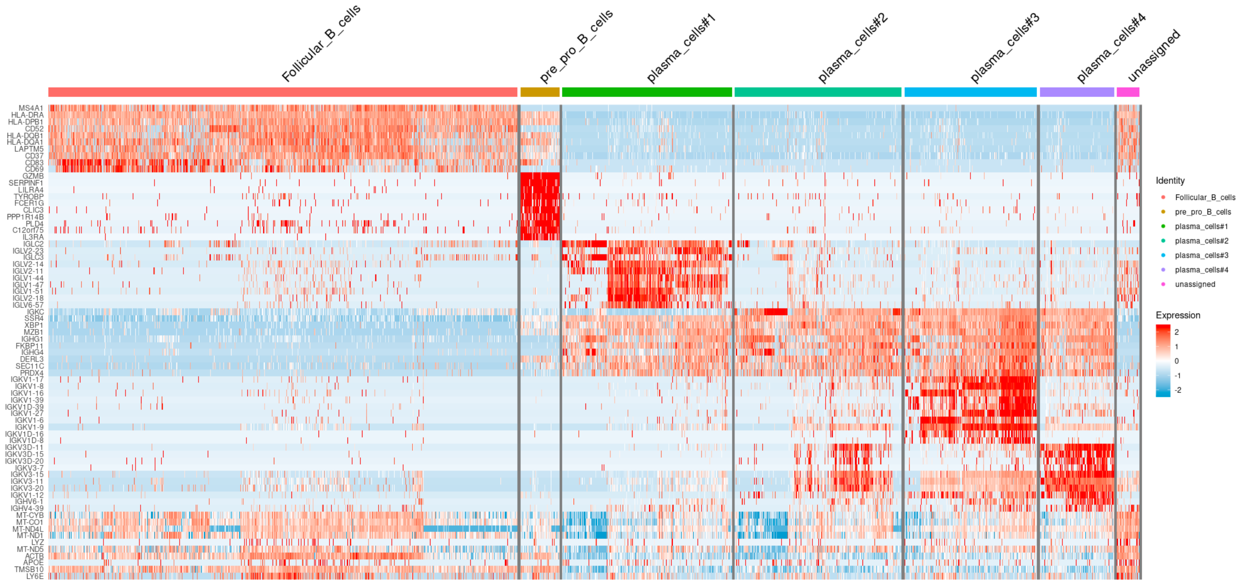


(e)

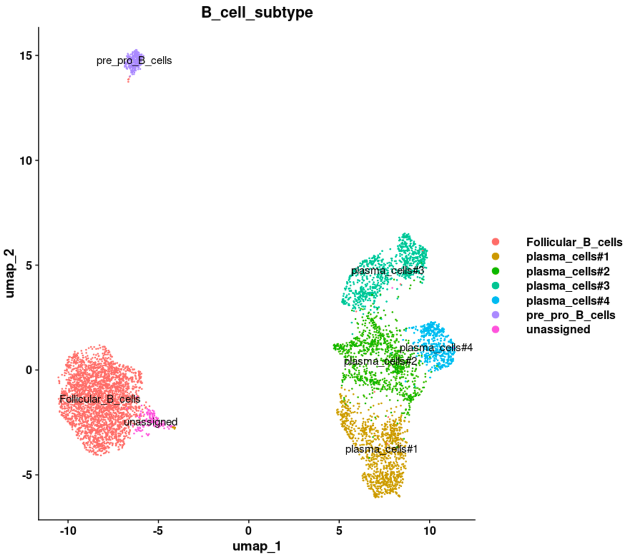


(f)

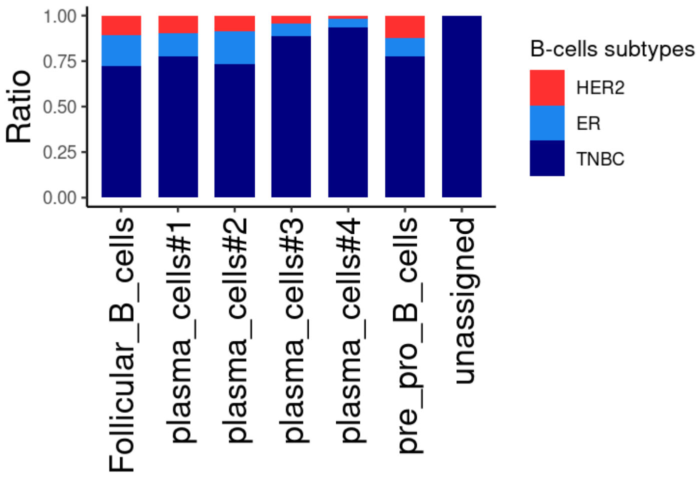

**Supplementary Figure S3**

(a) Feature plots illustrating the expression of markers genes used for identification of T-cells/ NKT cells subtypes.

(b-c) Feature plots illustrating the expression of markers genes used for identification of CAF subtypes, including matrix CAFs, collagen CAFs, interferon response CAFs and inflammatory CAFs

(d) Heatmap showing the transcriptional dynamics of tumor-associtaed B-cells in breast cancer

(e) UMAP illustrating the subtypes of tumour-associated B-cells in breast cancer

(f) Stacked bar chart showing the relative proportion of different types tumour-associated B-cells among different clinical subtypes of breast cancer.

### Supplementary Figure S4

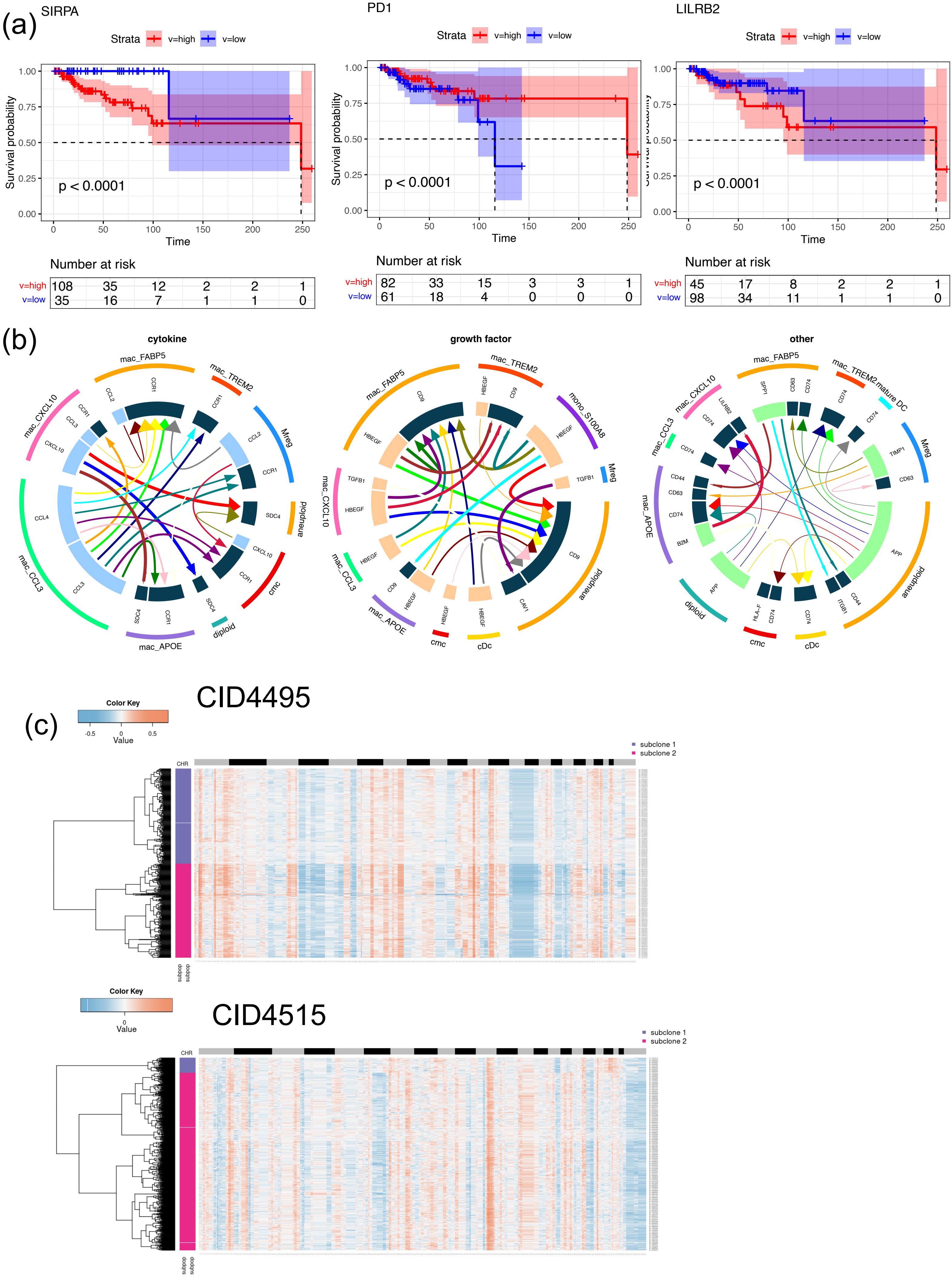

Supplementary Figure S4

(d)

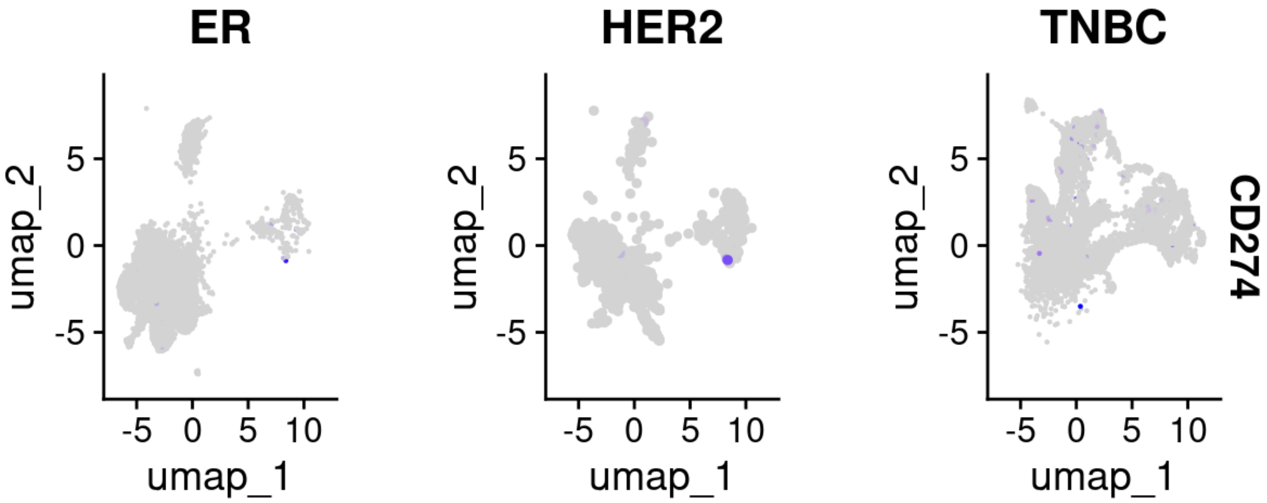

**Supplementary Figure S4**

(a) Kaplan-Meier survival analysis of the myeloid immune-checkpoint *LILRB2*. The dashed lines indicate the median survival time for the high-risk and low-risk cohorts. The 95% confidence interval bands for the high-risk and low-risk cohorts are shaded in low-opacity red and low-opacity cyan, respectively. The rectangle box below the curves contains count of individuals who are at risk over the given period.

(b) Chord diagrams illustrating the potential receptor-ligand interactions between myeloid subsets and aneuploid epithelial cells in breast cancer. The receptor-ligand interactions were classified as 'cytokine', 'growth factor' and 'other'.

(c) CNV analysis of the aneuploid epithelium-derived tumor cells from two representative TNBC patients (CID 4495 and CID 4515).

(d) UMAP illustrating the low expression level of PD-L1 (encoded by *CD274*) across the breast epithelial cells of different clinical subtypes

Supplementary Figure S5

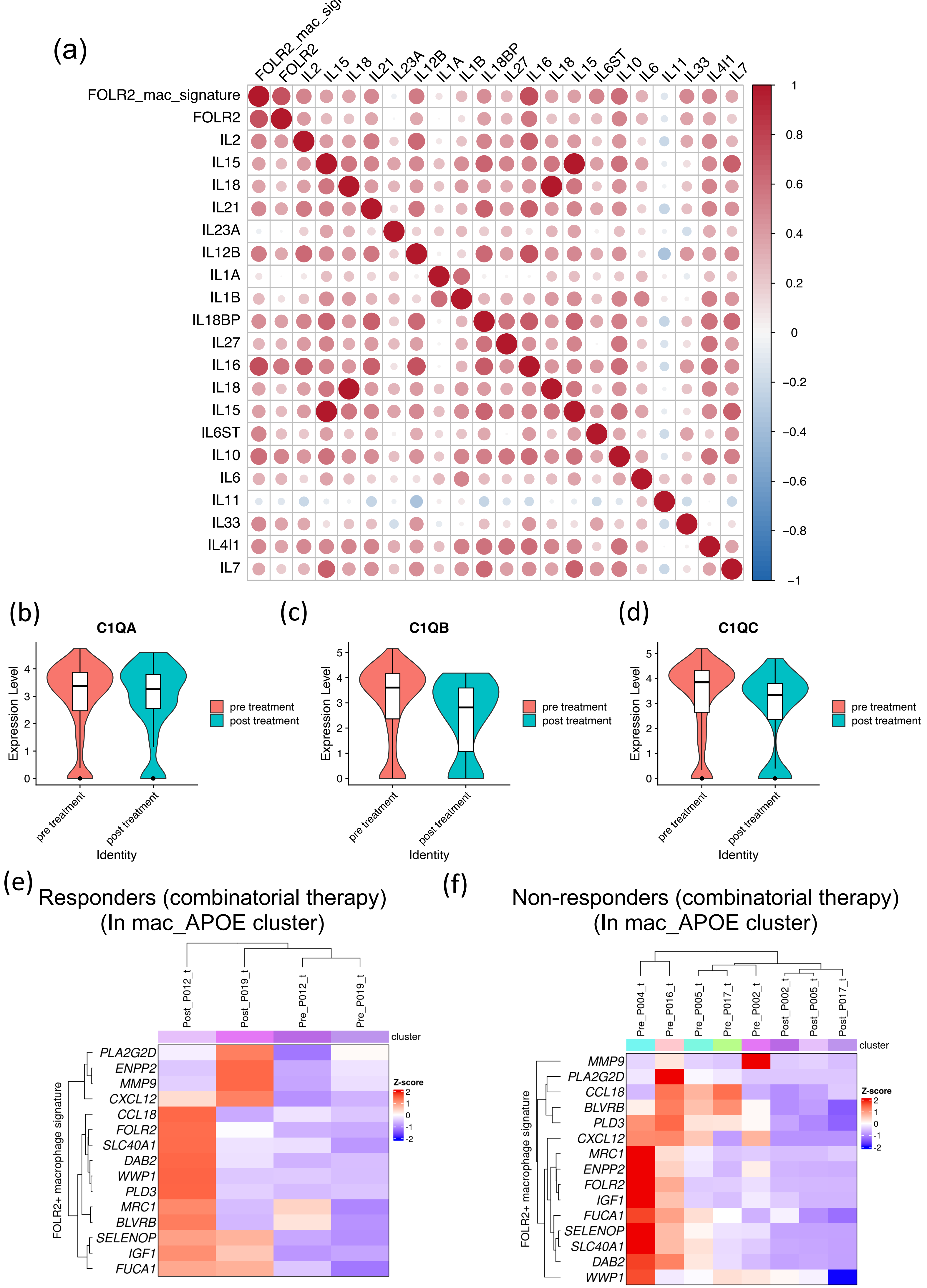

**Supplementary Figure S5**

(a) Bubble plots illustrating the spearman correlation between the expression of FOLR2_mac gene signature and interleukin genes. The correlation analysis was conducted using the bulk-RNA seq data of the BLBC patients in the TCGA-BRCA cohort.

(b-d) Violin Plots illustrating the expression of selected complement genes before and after combinatorial treatment

(e-f) Heatmap illustrating the expressions of FOLR2 signatures in mac_APOE cluster between different patients (either responders or non-responders) before and after combinatorial treatment
